# Gene co-expression analyses of health(span) across multiple species

**DOI:** 10.1101/2021.04.08.439030

**Authors:** Steffen Möller, Nadine Saul, Israel Barrantes, András Gézsi, Michael Walter, Péter Antal, Georg Fuellen

## Abstract

Health(span)-related gene clusters/modules were recently identified based on knowledge about the cross-species genetic basis of health, to interpret transcriptomic datasets describing health-related interventions. However, the cross-species comparison of health-related observations reveals a lot of heterogeneity, not least due to widely varying health(span) definitions and study designs, posing a challenge for the exploration of conserved healthspan modules and, specifically, their transfer across species.

To improve the identification and exploration of conserved/transferable healthspan modules, here we apply an established workflow based on gene co-expression network analyses employing GEO/ArrayExpress data for human and animal models, and perform a comprehensive meta-analysis of the resulting modules related to health(span), yielding a small set of health(span) candidate genes, backed by the literature.

For each experiment, WGCNA (weighted gene correlation network analysis) was thus used to infer modules of genes which correlate in their expression with a “health phenotype score” and to determine the most-connected (hub) genes for each such module, and their interactions. After mapping these hub genes to their human orthologs, 12 health(span) genes were identified in at least two species (ACTN3, ANK1, MRPL18, MYL1, PAXIP1, PPP1CA, SCN3B, SDCBP, SKIV2L, TUBG1, TYROBP, WIPF1), for which enrichment analysis by g:profiler finds an association with actin filament-based movement and associated organelles as well as muscular structures.

We conclude that a meta-study of hub genes from co-expression network analyses for the complex phenotype health(span), across multiple species, can yield molecular-mechanistic insights and can direct experimentalists to further investigate the contribution of individual genes and their interactions to health(span).

## Introduction

Health and healthspan are gaining acceptance as central concepts in medicine, with a focus on (multi-)morbidity, aiming to delay the onset of disease and dysfunction for as long as possible. Health is difficult to describe and has different meanings to different people. Aging, and the deterioration of health that comes with it, affects nearly all species. But tissues that enable the systematic study of the underlying molecular processes are more easily available for animal models, especially for invertebrates, coming with further advantages such as controlled genetics and environments, and a much shorter lifespan. Thus, aging and healthspan are frequently studied in animal models.

To support aging research, many databases are now available (Tacutu *et al.*, 2018). Gene expression profiles across tissues of aging mice were already presented, e.g., by the AGEMAP (Zahn *et al.*, 2007) project in 2007 and recently by the Aging Atlas Consortium (2020), but there is a lack of such data for healthspan. Adding the dimension of health may amend the identification of molecular markers for aging and further support the identification of health-modulatory compounds (Dönertaş *et al.*, 2018).

An increasing number of transcriptomic data sets that can be used to compare young and old individuals are available on public repositories. The concept to derive aging-associated patterns from transcriptome repositories across species (de Magalhães *et al.*, 2009) already led to central elements of aging-related knowledge bases (Tacutu *et al.*, 2012 and 2018). Comprehensive analyses of transcriptome repositories were also expanded towards diseases in the context of aging (van Dam *et al.*, 2012). Yet, as for expression profiles *per se*, there is a lack of gene expression co-regulation analyses across species with a focus on health(span). A major challenge for polygenic phenotypes in general is the heterogeneity of the underlying gene regulatory landscape (Kotlyar *et al.*, 2019), impeding the use of network-based methods for post-processing, i.e., smoothing, aggregating, and unifying, transcriptomic results (Leiserson *et al.*, 2013; Cowen *et al.*, 2017). However, the power of the cross-species derivation of conserved co-regulation modules is becoming apparent, see, e.g., the CoCoCoNet database (Lee *et al.*, 2020).

For prominent cellular characteristics of aging, such as cellular senescence, Avelar and coworkers (2020) demonstrated how to integrate static data from public databases with insights from gene co-expression (https://coxpresdb.jp/, Obayashi *et al.*, 2019). Attempts were also made to use known gene/protein interactions to describe age-induced expression profiles (Faisal and Milenković, 2014). The integration of co-expression data, also across species, could similarly be performed with GeneFriends (van Dam *et al.*, 2015, for human and mouse) for RNA-seq or, for microarray data also with MIM (Adler *et al.*, 2009). The latter also provides provenance information, i.e. the experimental context in which the correlation was found, to plan follow-up experiments.

We recently proposed an operational definition of health (Fuellen *et al.*, 2019) and suggested that it may be applied across species. We then collected data on molecular contributions to health (Möller *et al.*, 2020), with a focus on genetics. With the support of GeneMania (Franz *et al.*, 2018) and the associated tool AutoAnnotate (Kucera *et al.*, 2016) we then constructed a map of network modules by clustering a functional interaction network of the genes implicated in health. Naturally, aging and health are complex phenotypes for which we still lack the means to single-out and investigate the contribution of individual genes. Any detailed analysis is therefore expected to dissect a list of health-associated genes into gene sets that, in turn, can be understood as parts of the whole (that is health), and these parts are distributed across diseases & dysfunctions, tissues & organs, and species. The idea of identifying health-associated molecular patterns is at the root of molecular health research. Our efforts strived for a consensus across the species barrier of worm (*Caenorhabditis elegans*) and humans and we investigated the transfer of findings from worm as a short-lived animal model of health to humans. A consensus in network modules of worm and human was thus determined (Möller *et al.*, 2020), but it was small in relation to the much larger functional interaction networks that were the starting point for each species. However, functional interaction databases, upon which GeneMania is based, are woefully incomplete. Also, these databases do not usually consider the specific biological context of an interaction, but instead merge interaction data from very heterogeneous sets of experiments (Magger *et al.*, 2012; Kotlyar *et al.*, 2019).

To harness the power of diverse transcriptomic experiments in the context of health(span), here we present a WGCNA-based meta-study for the exploration and characterization of health(span) related modules. WGCNA co-expression analyses have recently been used in aging research (Li *et al.*, 2019) to identify differences in old vs young and gene expression asymmetries *in the brain* that develop over time. In our study we integrated a very diverse set of health(span) expression data across species from many different tissues. We manually derived a scoring for all the transcriptome samples we consider, based on a score combining quantitative and qualitative factors that the authors of the experiments provided and refer to it as their “health phenotype score”. WGCNA was found to be a competitive tool to find network modules reflecting such kinds of scores (van Dam *et al.*, 2017). Across tissues (or cell lines) and multiple species, this allows the filtering for health-associated modules generated by the WGCNA correlation analysis and thus, the meta-study of health-associated most-connected genes (hubs) and of their interactions, as presented here. We also collected the evidence for the implication of these genes in health(span) from the literature.

## Methods

All sets of transcriptomics experiments in the Gene Expression Omnibus (GEO, Clough and Barrett, 2016) and ArrayExpress (Athar *et al.*, 2019) databases that mention “healthspan” in the title or the description were included, if they featured more than 6 samples and a scale-free network could be derived from their correlation matrix (for the latter, see below). Experiments performed on *C. elegans* were added when these were alternatively annotated with the term “health”, to increase the number of datasets for the worm, since ‘healthspan and "Caenorhabditis elegans"’ only finds the single entry E-GEOD-54853. We did not include *non*-worm experiments with “health” in the title or the description, since the number of matches (specifically for human) turned out to be excessively large.

Each experiment’s metadata was inspected to manually derive a score designed to reflect the health status of the individual(s) from which the sample(s) were taken, unless such a score was already given. This “health phenotype score” was manually tailored for each experiment by a custom formula that takes the experiment’s factor annotation as an input and thus consistently annotates each sample. This can be inspected in the ‘Data_parameters’ folder (see Availability). Log-transformation of expression levels was performed if not already performed for the data we retrieved. Table 1 describes the experimental data and metadata which form the input to the following analyses.

**Table 1:**
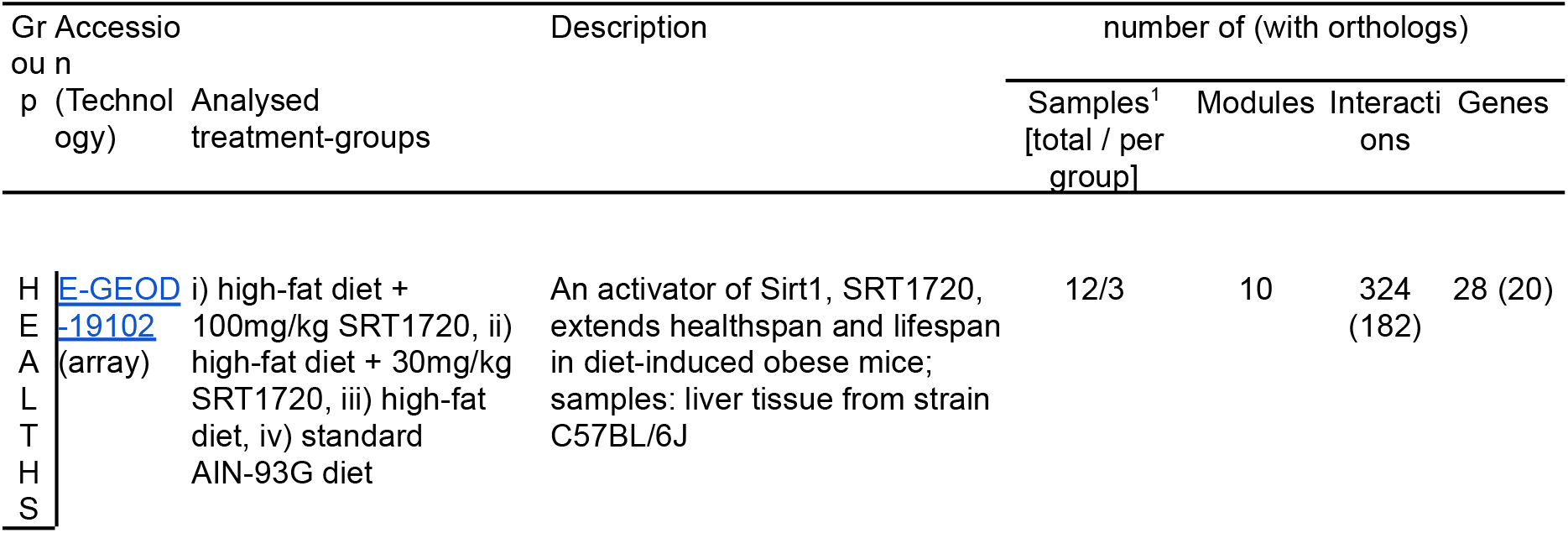
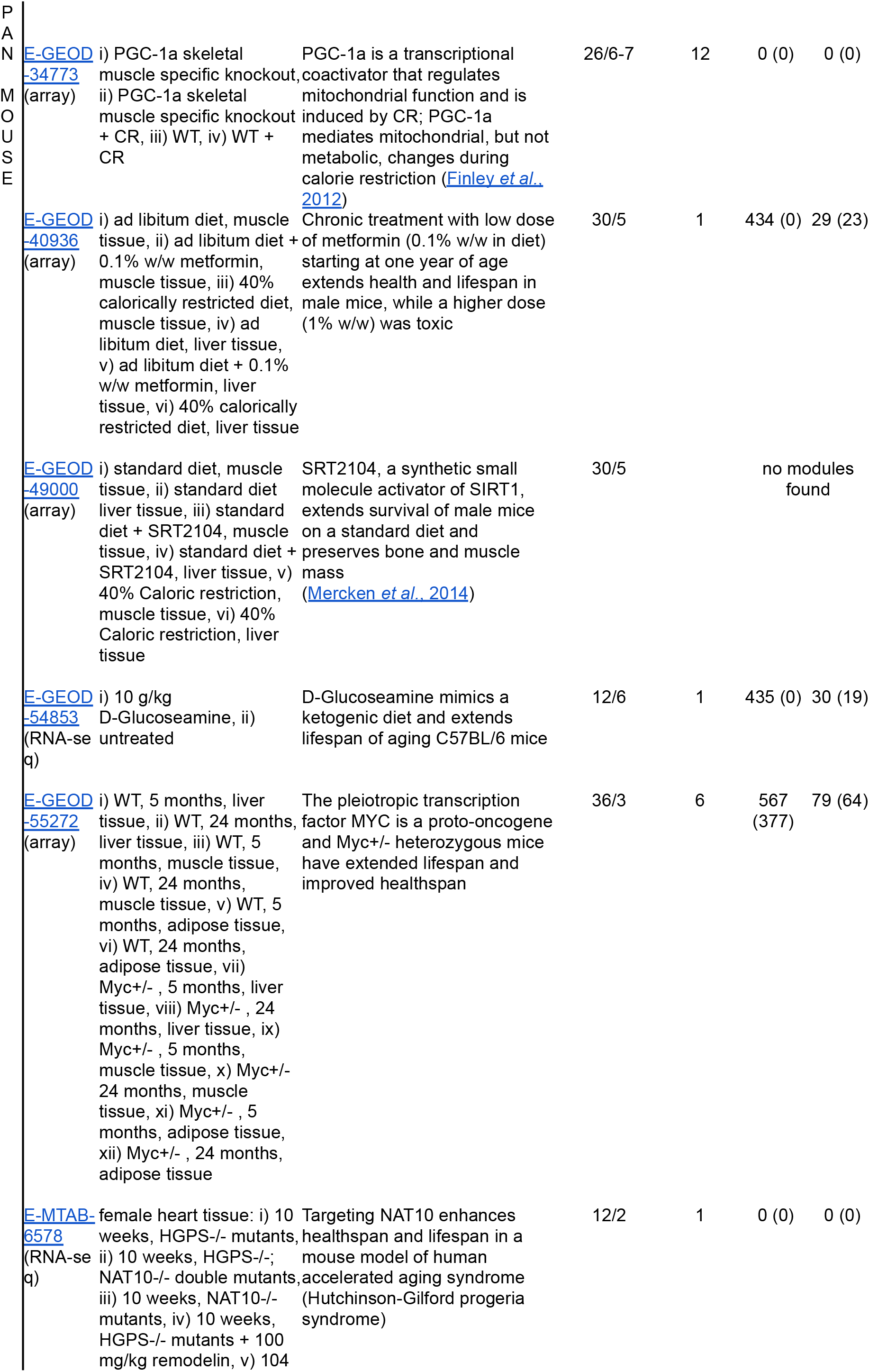
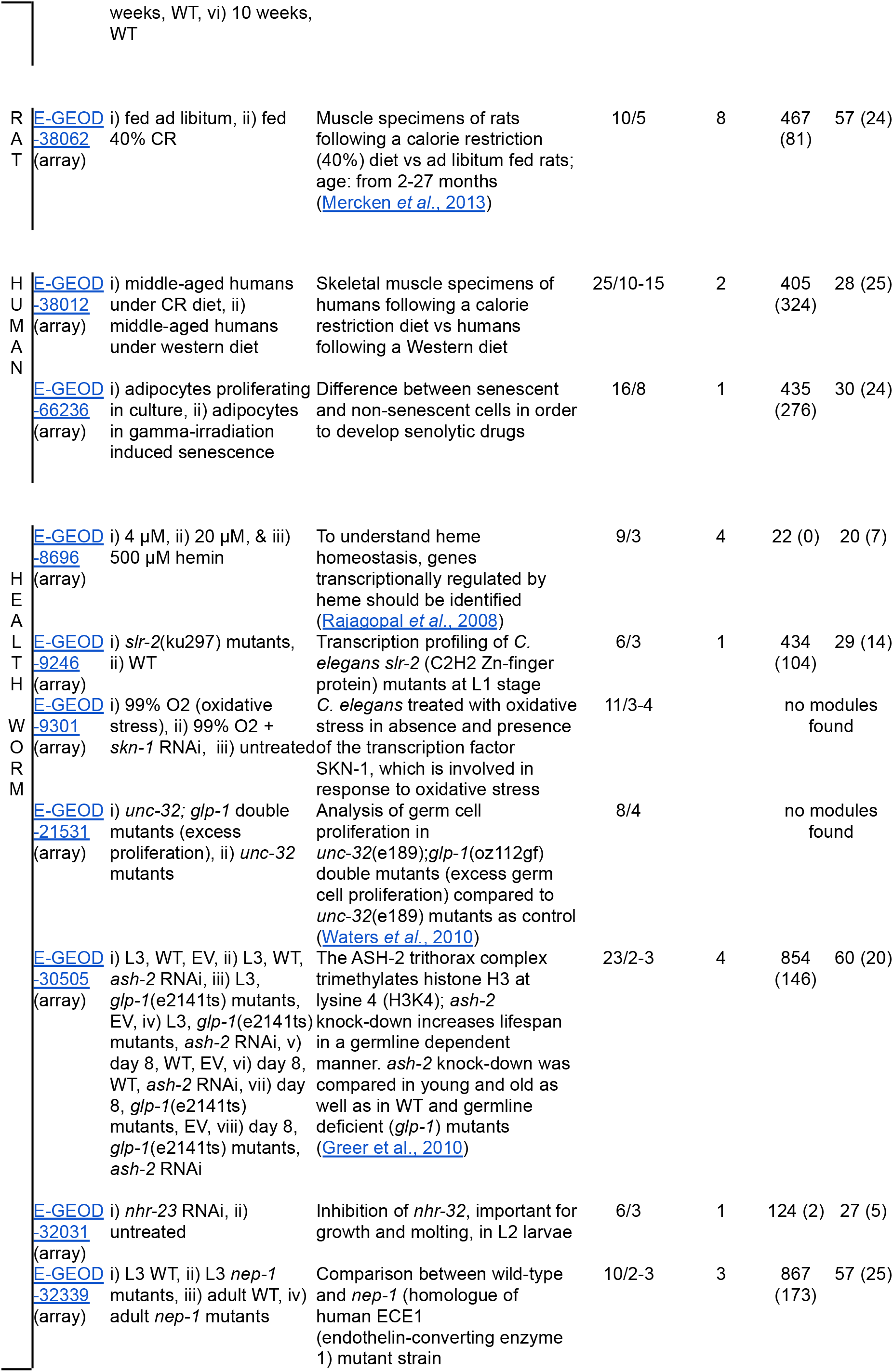
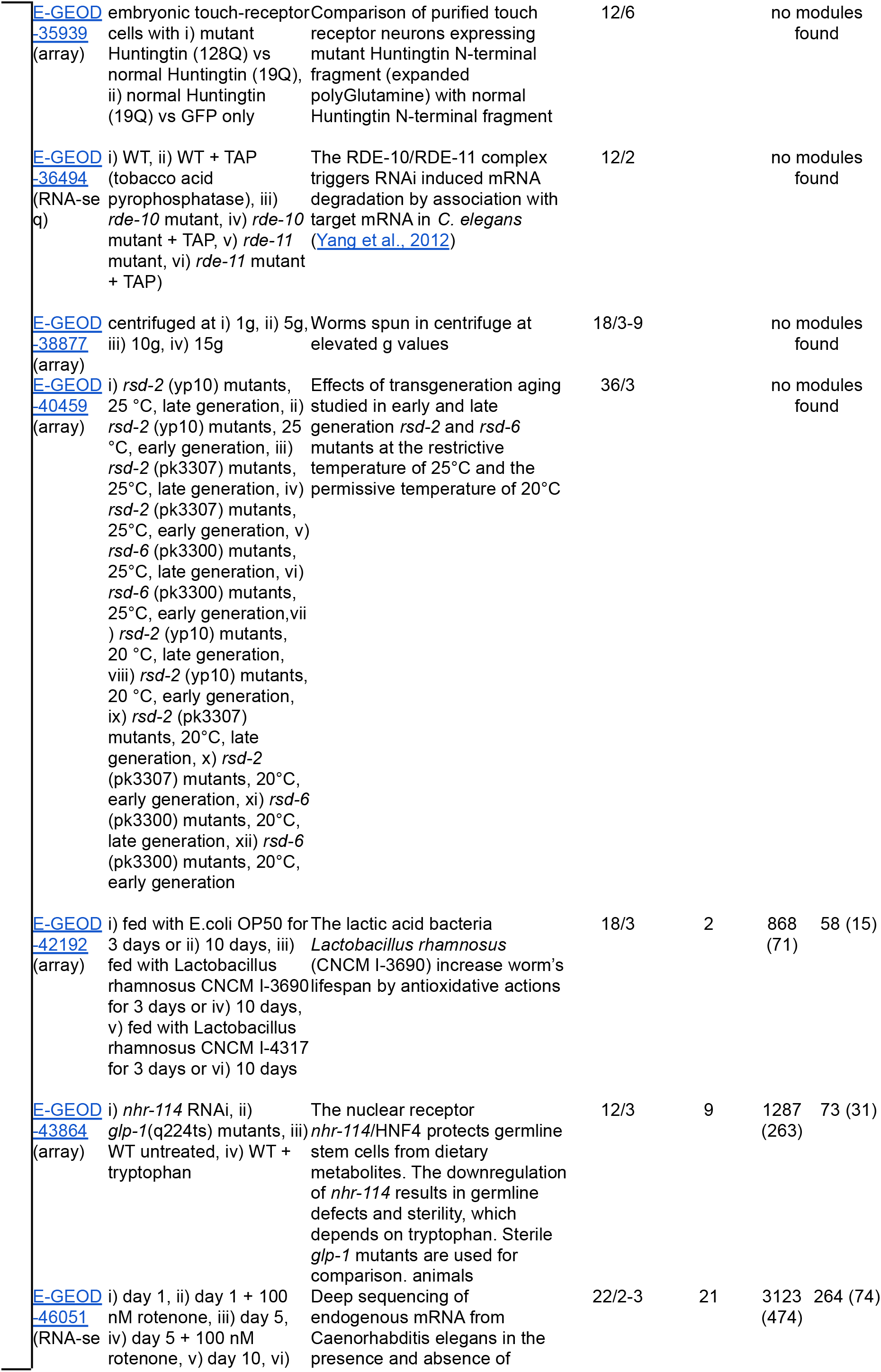
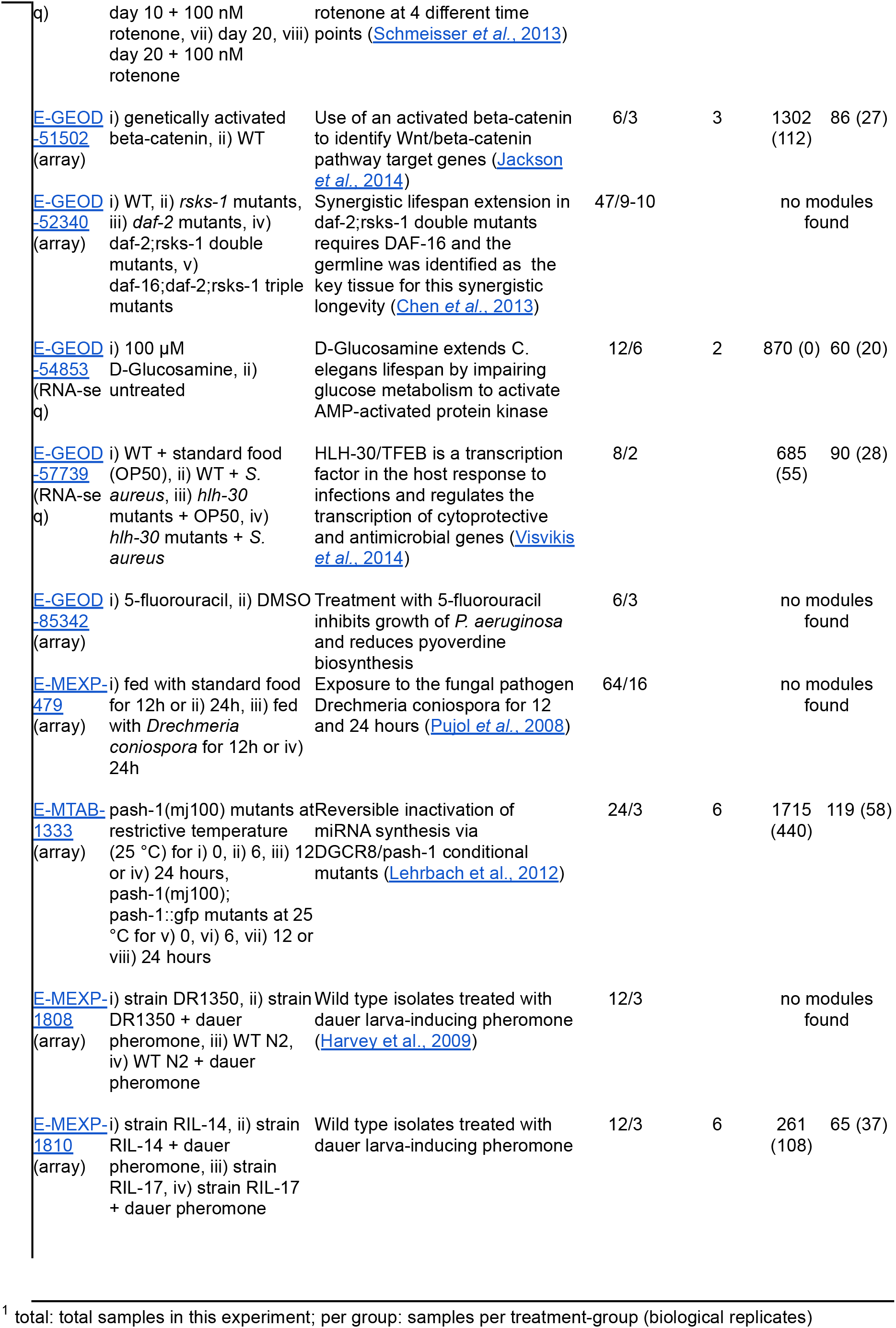
List of ArrayExpress/GEO files used as input in our study. *This table provides an overview of the transcriptomics experiments that were retrieved for this study. Each experiment was processed by a regular WGCNA workflow with unsigned correlation. Interactions were collected for the 30 most connected (hub) genes in each module. The column Modules lists the number of modules found for the experiment that feature an eigengene that correlates (with P<0.05) with the samples’ health phenotype score. Within each module, only interactions with an adjacency value above the 95^th^ percentile of an experiment were considered. The rightmost column lists the number of different hub genes that are paired in any of these interactions. Numbers in parentheses give the number of genes/interactions that could be mapped to ortholog genes in the human; for human data, the number of orthologs in worm are shown. For an interaction, both of the paired genes need to have orthologs assigned; otherwise they were not considered for the count*.

### RNA-seq data re-analysis

Gene expression levels were typically not available for the RNA-seq data. Therefore, the RNA-seq datasets were all reanalyzed based on the raw data by the following protocol. All target RNA-seq datasets were retrieved from the European Nucleotide Archive (Leinonen *et al.*, 2011), and the corresponding FASTQ files were filtered for Illumina adapters, phage PhiX sequences and quality (Phred score over 25) using BBTools version 38.49 (Bushnell *et al.*, 2017). Gene expression was then quantified for each RNA-seq run. To this end, the filtered outputs were mapped against the corresponding target genomes from the Ensembl database release 98 (Yates *et al.*, 2019), using the STAR program version 2.7.3a (Dobin *et al.*, 2013). This program also enabled us to assign uniquely mapped reads to individual genes from the short read alignments. Finally, the mapped read counts were normalized as transcripts per million (Li *et al.*, 2010).

### Network analysis

For each experiment, gene interactions are derived from their pairwise Pearson-correlation of gene expression across samples with the WGCNA (Langfelder and Horvath, 2008; Zhao *et al.*, 2010) R package. The WGCNA analysis was performed for undirected interactions. Parameters were set as instructed by the WGCNA standard protocol, as follows. For every experiment the *cutHeight* was manually set to remove outliers and the exponent/power was manually determined to ensure that the network is a scale-free network (see below). An experiment is skipped if that is not possible and then marked with “no modules found” in Table 1. For RNA-seq, prior to the removal of outliers, low-count genes were removed by a manual setting of the parameter *cutHeight* so that the separation of the samples reflects their phenotypes and could no longer be improved, based on the clustering of the genes by expression data with the R function *hclust* as performed as part of the WGCNA protocol.

In an attempt to make the correlation networks of different experiments more similar to each other with respect to the number of connections that may be expected for each gene, we adhered to the WGCNA protocol that proposes to apply an experiment-specific exponent to the correlation coefficients (WGCNA calls it “power”, which it is, but not in the context of the power law mentioned below) to strengthen the differences in the correlation data. Therefore, this power is chosen, for each experiment, just large enough so that in the derived correlation network, the fraction of genes that have *k*-many interactions with other genes is proportional to k^−γ^ with γ being a small positive parameter. Networks with that property are called scale-free; the parameter γ describes how quickly this fraction gets smaller when the number of connections increases. Genes with a high number of connections are rare in scale-free networks, but they exist, and these “hub” genes are considered highly influential on the expression levels of genes in that module. Further, we filtered for modules that are associated with the health(span) phenotype (see next paragraph), and the hub genes are likely to also have a strong effect on this phenotype.

Only network modules whose WGCNA eigengene correlated with the “phenotypic health score” (p-value < 0.05) were retained further. Then, the 30 genes (see the WGCNA tutorial https://horvath.genetics.ucla.edu/html/CoexpressionNetwork/Rpackages/WGCNA/Tutorials/) most connected in a module according to the WGCNA *softConnectivity* function were considered for subsequent consensus analyses, and called “hub” genes hereafter. Besides the modules, output of the WGCNA workflow is the topological overlap matrix with a quantitative description (termed *adjacency*) of all interactions between any pair of genes of an experiment. For each experiment, we determined a threshold at the 95% quantile of all the adjacency values. Only gene interactions with an adjacency above that experiment-wide threshold contribute to our analysis of interactions of genes in the health-associated modules. For the 30 hub genes, all pairwise interactions above that experiment-wide 95% quantile were thus exported. These interactions were then subjected to a pan-module search for consensus genes and consensus interactions, also across species by considering orthologs, presented in Tables 2 and 4. Orthologs were determined based on Ensembl version 101 (Yates *et al.*, 2019).

**Table 2a:**
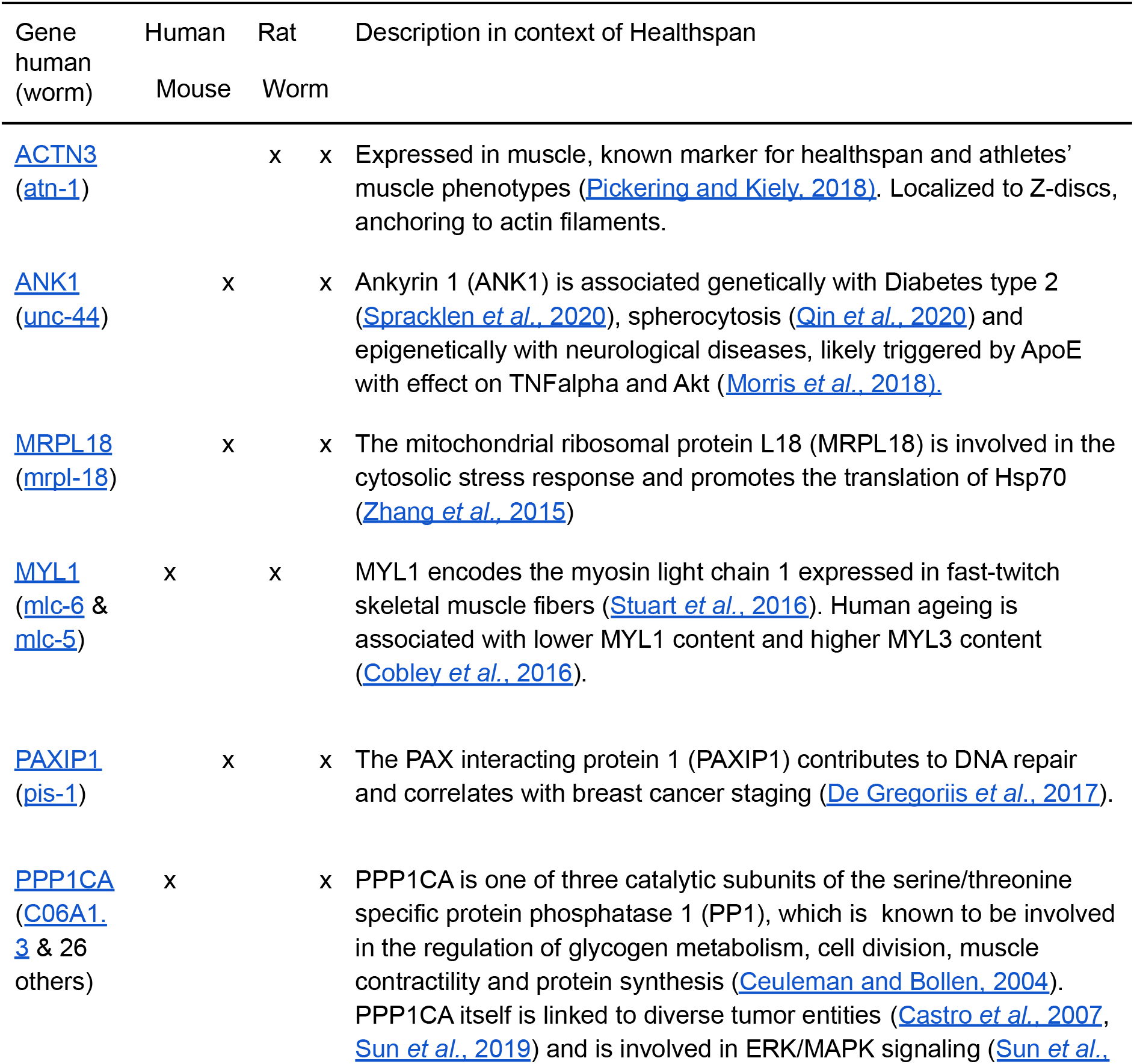
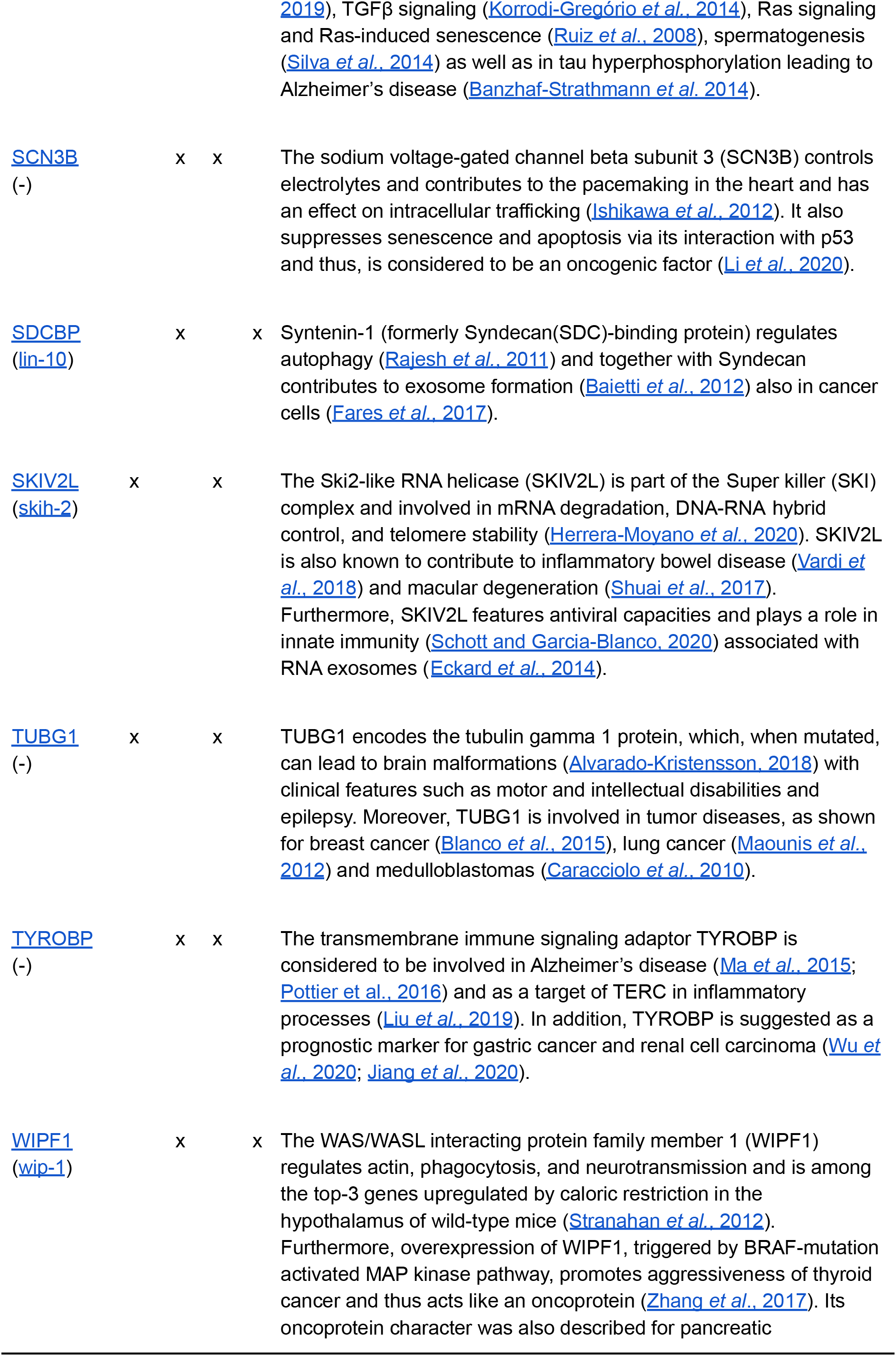
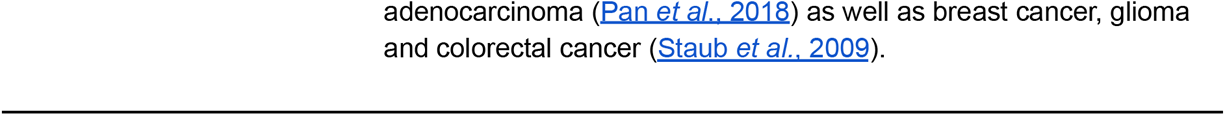
Hub genes in health-associated WGCNA network modules, found in at least two species. *Orthologs were mapped to the human gene name using Ensembl. The human gene names also correspond to the names in mouse and rat, whereas the names of the orthologs in worms based on the Ensembl database are given in brackets*.

Further, for each health-associated module, the 30 genes correlating the strongest with the module’s eigengene (reflecting average module behavior, also called “module membership” in WGCNA) were retrieved. Those found in at least two species are presented in Table 3. The correlation is taken in relation to the eigengene, and not in relation to the health phenotype score. Either would be fine for a ranking of the hub genes within a module, and the ranking is expected to be identical for the genes most central to a module. However, our particular interest was to abstract from the phenotypes of the experiment and thus utilize the WGCNA-performed modularization to influence the ranking. This is assumed to be particularly useful for experiments with multiple health-associated modules, each of which we expect to focus on a different aspect of health and for which hence also the genes should be ranked differently, to help analyzing that particular module most appropriately, and without any particularities pertaining to the health phenotype score. Multiple probesets describing the same gene, or its splice variants, were not distinguished and mapped to the same human gene, resulting in genes interacting with themselves. Such self-interactions were removed.

**Table 3a:**
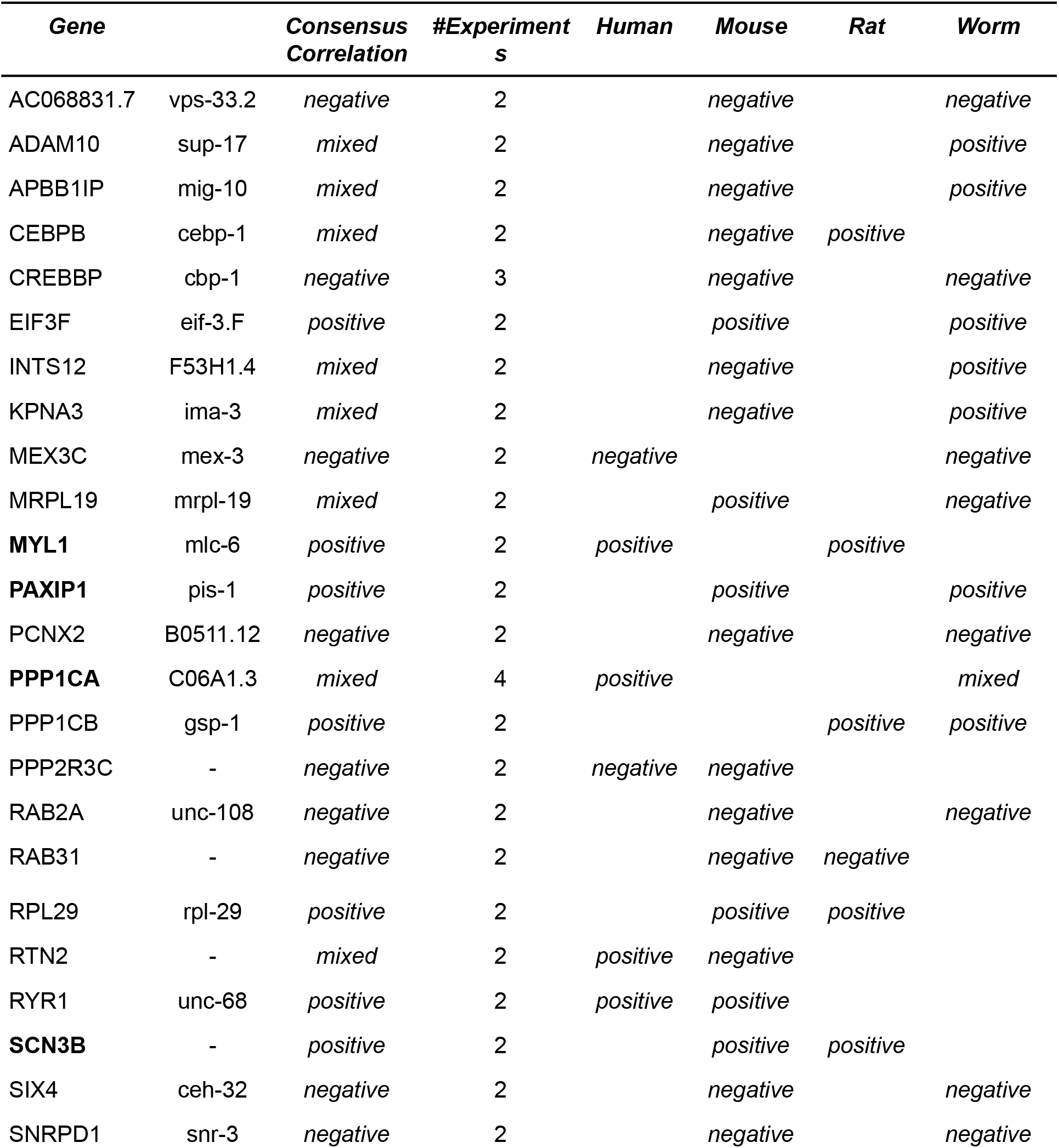
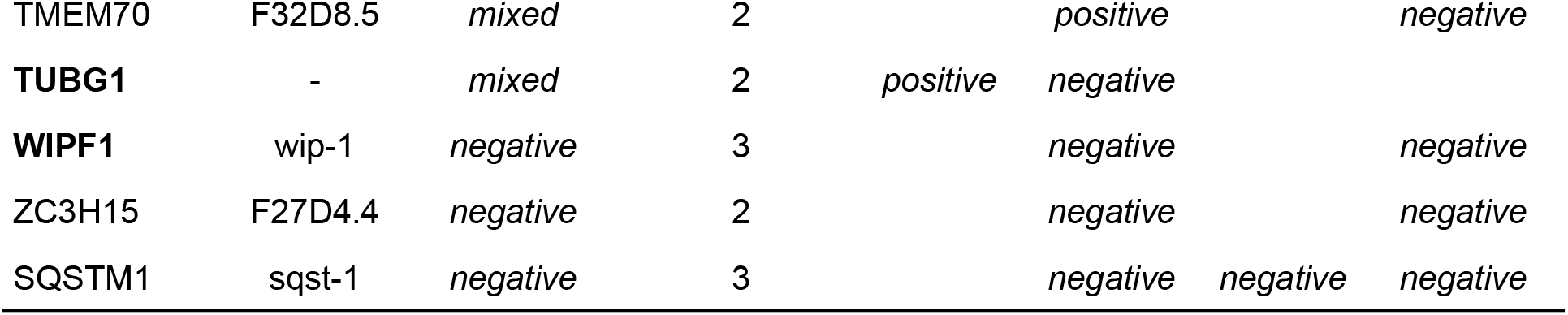
Genes correlating the strongest with the module’s eigengene (quantifying module membership) in at least two species. *Genes in this table are among the top-30 of the module membership and found in experiments of at least two species. The gene name is marked in bold if that gene was listed as a hub gene in Table 2. The column “Consensus Correlation” flags “positive” (or “negative”) to refer to an observed positive (or negative) correlation with the “health phenotype score” when the gene is upregulated. “mixed” indicates that the experiments did not yield a consensus direction of correlation. Supplement Table 1 extends this list to all genes that appear in the top 30 of modules of two or more experiments. The “#Experiments” column indicates the number of experiments with a module for which the gene was identified as a member*.

### Network figures

The network figures were created with the R igraph package. A spring-embedding layout was chosen for the plots and manually refined. Figure 2 shows an overview on all hub genes from Table 2, their direct interactions and genes found in any species that connect to at least two hub genes. Additionally, Figures 2a-d in the Supplement were prepared separately for each species, i.e. they show only interactions from modules that WGCNA identified for an experiment based on samples from that species. Input to these supplement figures are the hub genes from Table 2 and all genes that are reachable from the hub genes which are no more than two transitions away. All interactions between the selected (reachable) genes were also added. The resulting graphs were simplified with the igraph minimum spanning tree implementation that maintains the connectivity of the graph but removes all redundant paths between genes. The spanning tree retains the stronger of two alternative paths between genes. A gene connected to a hub gene with a low adjacency value will thus lose that direct link if it is correlating strongly with another gene that has a strong correlation with that hub gene. Hub genes were determined from within WGCNA considering all interactions, not only the ones above the 95th percentile. Hub genes that are strongly connected for experiments in one species may not be equally dominating in another species. This and the competitive effect on directly connected hub genes (one cross-species, the other only observed for one species) imposed by the spanning tree give the impression that the cross-species consensus hub genes are marginalized in Supplemental Figures 2a-d, albeit these graphs are seeded from the consensus hub genes and their interactions.

**Figure 1:**
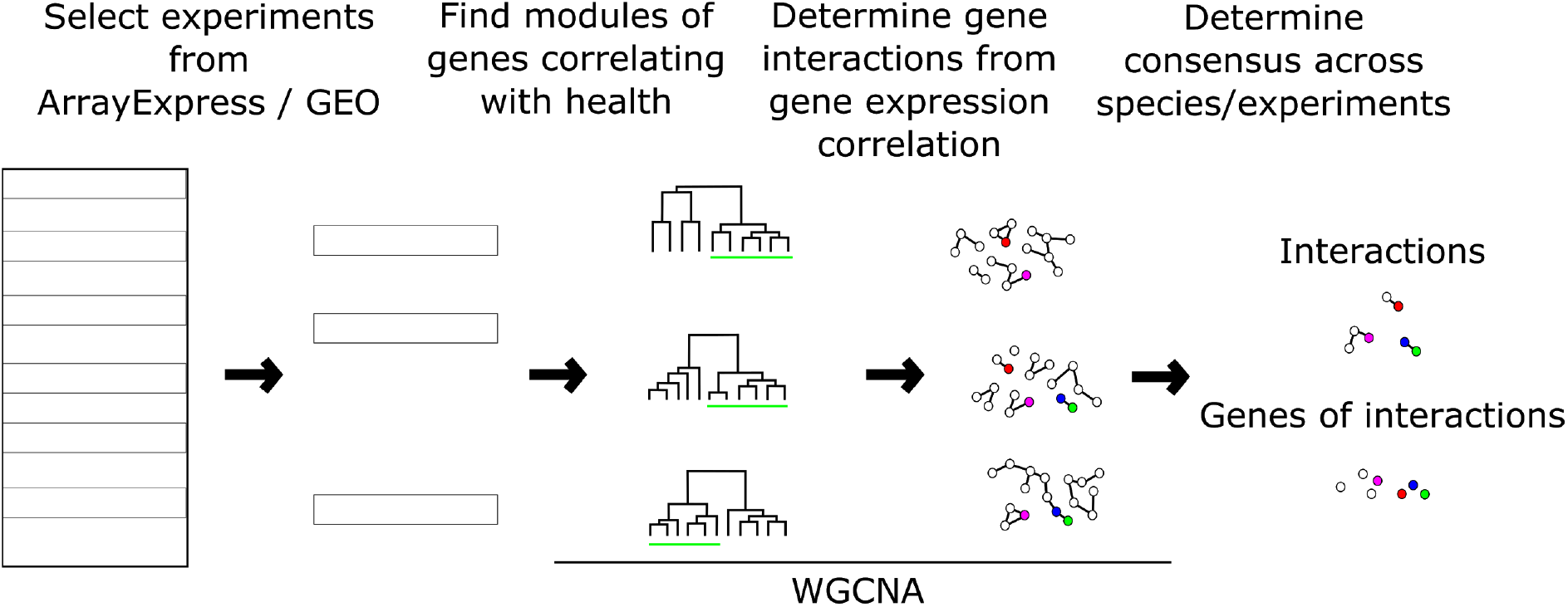
Workflow to determine cross-species consensus gene correlation networks, and subsequent analyses. WGCNA is applied independently for each selected experiment in ArrayExpress/GEO, defining modules and gene interactions. Gene interactions are filtered by experiment-specific thresholds. For each module, hub genes are retrieved and those with an ortholog found as a hub gene in another species are reported in Table 2. For each module, Table 3 lists the genes that correlate the most with its “eigengene”, i.e. that best represent the module’s expression pattern across samples.

**Figure 2:**
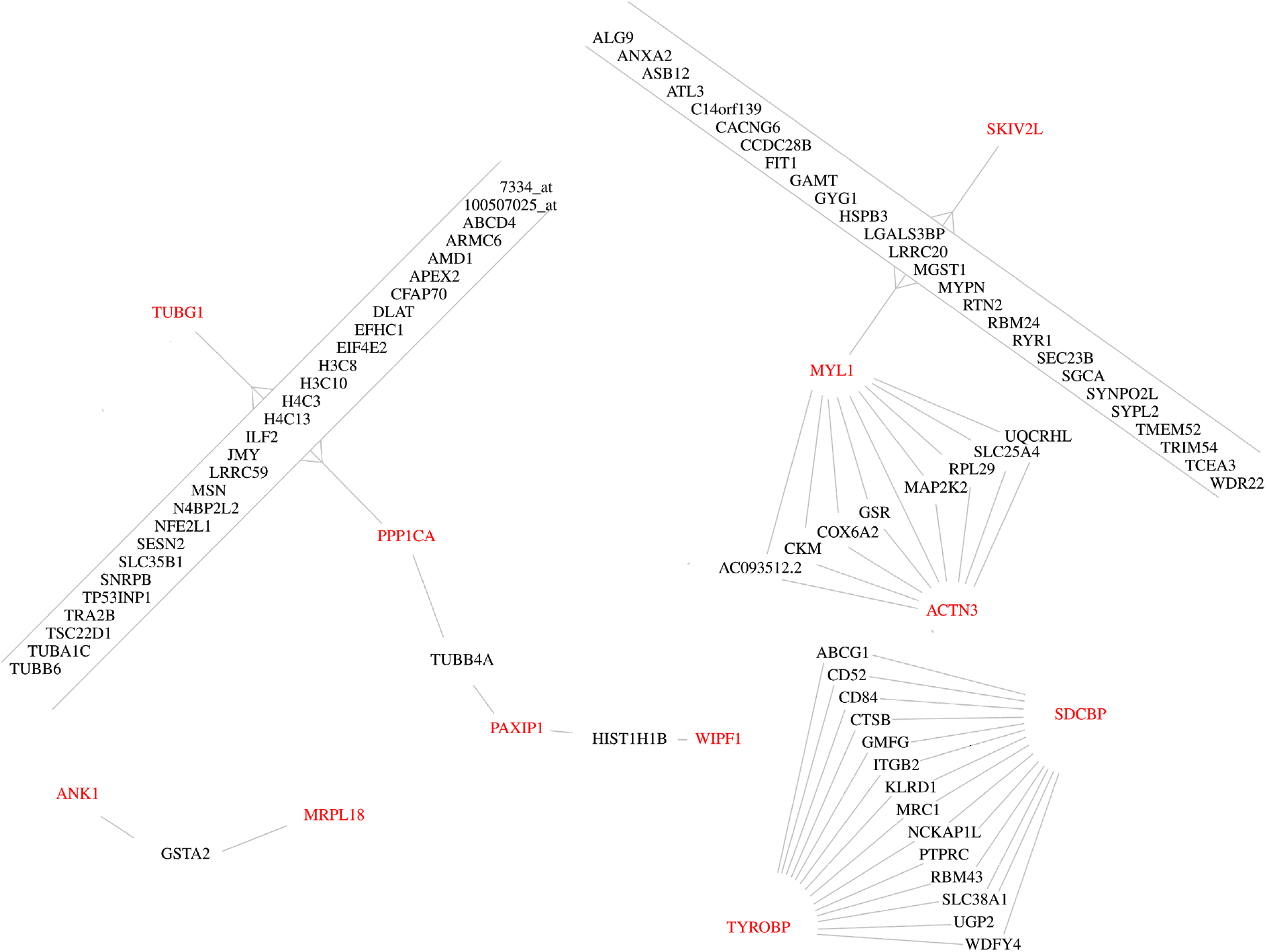
Cross-species conserved hub genes observed in health(span)-associated WGCNA modules, and genes that connect these hub genes. Connections are interactions taken from the WGCNA adjacency matrix if the adjacency is above the 95^th^ percentile of all interactions of that experiment and if for that experiment the interaction is in a health(span)-associated module. The only direct interaction between hub genes is between MYL1 and ACTN3.

## Results

We analysed all experiments listed in Table 1 with WGCNA. This analysis provided a modularization by an expression-based clustering of genes and allowed to describe the association of each module with the “health phenotype score”. WGCNA also quantified the strength of gene correlations and determined hub genes for each module. We identified 12 genes (Table 2) that are among the 30 hub genes in health(span)-associated modules from at least two species. In total (Supplemental Table 1), 658 different genes were found among these top-30 hub genes of all modules as determined by WGCNA. An interaction network of the genes from Table 1, based on correlation of gene expression, is presented in Figure 2.

To prioritize the cross-species hub genes of Table 2, we also looked at the module membership of all genes for each module. The genes most correlating with the module’s eigengene are reported and, analogous to Table 2, the genes that are found in multiple species were determined and listed in Table 3. This table further indicates whether a gene’s change in expression is positively or negatively correlated with the eigengene of the WGCNA module to which it belongs, which in turn may be positively or negatively correlated with the health(span) phenotype. Supplement Table 1 shows the raw data that were used to construct Table 3. To allow for a direct comparison of the genes’ correlation with health(span), not quantitatively but in terms of direction (that is, up- or downregulation in relation to the health phenotype score), Table 3 presents a gene’s inverted direction if the gene’s module is already negatively correlated with the health(span) phenotype. The “Consensus Correlation” presents the direction that all experiments are in agreement with or “mixed” if the experiments differ in terms of their correlation with the health phenotype score. This information can be calculated for all genes, which we consider to help interpreting a module. For each module, the data for the 30 genes correlating the strongest with the module’s eigengene are therefore provided in the supplement (Supplement Table 1).

The intersection of Tables 2 (hub genes) and 3 (genes correlating with the health phenotype score) points to a subset of genes that are considered both influential and directly associated with health, i.e. MYL1, PAXIP1, PPP1CA, SCN3B, TUBG1 and WIPF1. The enrichment by g:profiler for the genes of Table 2 are shown in Figure 3. Supplement Figure 1a shows an enrichment analysis for the intersection of Tables 2 and 3 which is matching closely the enrichments in Figure 3, except that it does not feature the terms associated with muscle.

Supplement Figure 1b shows the enrichment for all genes in Table 3. The latter is the least robust since the enriched terms do not cover a large fraction of the genes as for the other enrichment analyses.

**Figure 3:**
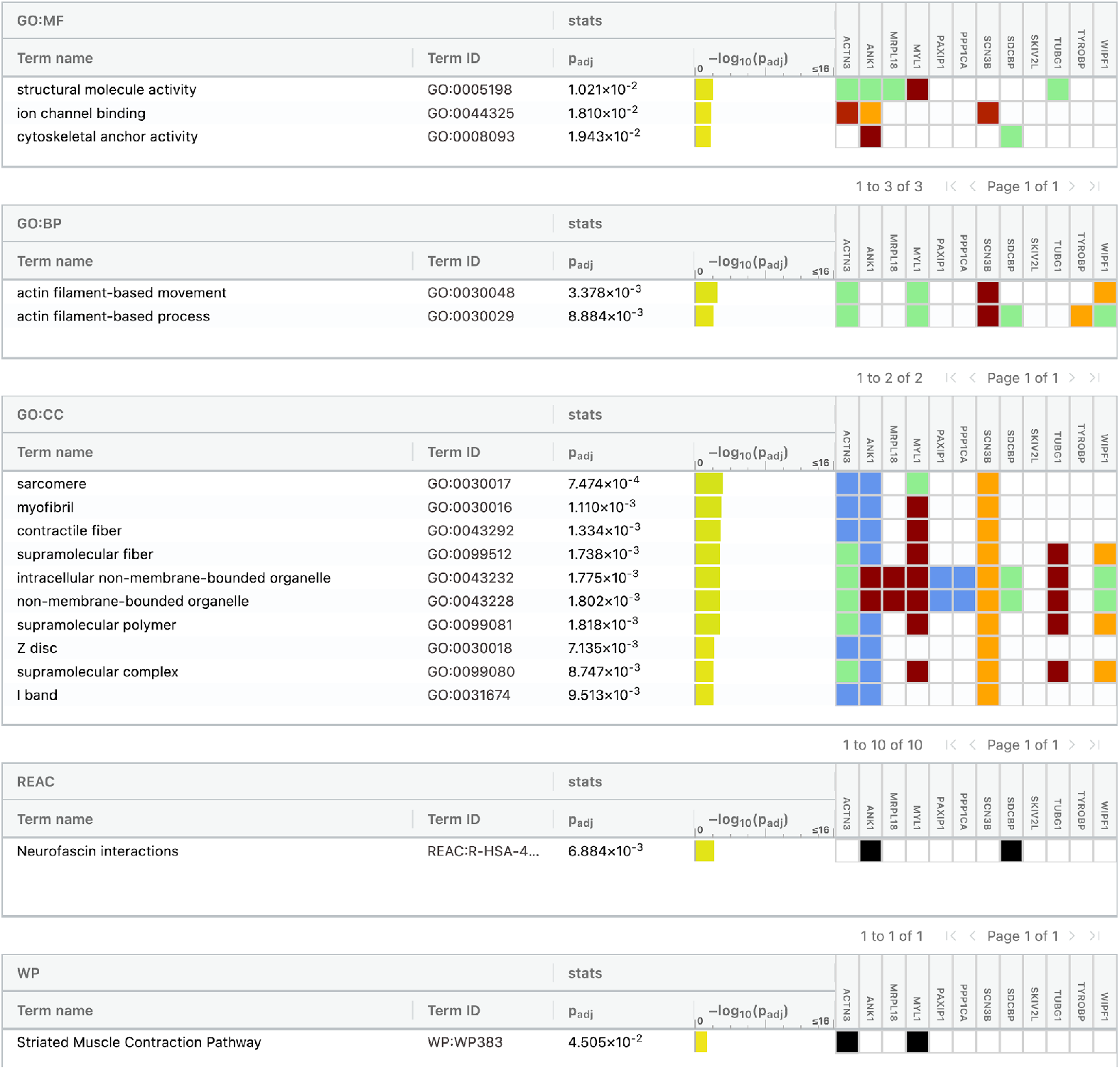
Gene set enrichment analysis of cross-species hub genes for health(span) with g:profiler. Input are genes from Table 2 that are observed in healthspan-associated modules of multiple species. Terms with a low coverage of genes are not suitable to describe the selection as a whole but may still direct the interpretation of parts of the network where these genes are connected.

## Discussion

### Method

The onset of this investigation were all experiments in GEO/ArrayExpress that mention “healthspan” in their description (or “health” or “healthspan” in case of worm). For each experiment, from the descriptions that are provided for the samples in the database, a “health phenotype score” was derived. A gene expression correlation analysis with WGCNA yielded a gene coexpression network for each experiment as a set of modules of genes that correlate with the health(span) phenotype. We were interested in genes that are most connected, i.e. hub genes, for each module, and in their interactions as described by the WGCNA network. The correlation of genes with the module eigengene (Table 3), to predict a positive or negative association with health in the molecular context of that module, was only of secondary interest to us.

In this analysis, we focussed on common observations across two or more species and a variety of health-related phenotypes, including the reaction to drugs that extend healthspan (Table 1). The first steps of our analysis with WGCNA identified modules directly from the expression data, i.e. without inspecting a phenotype; the selection of health(span)-associated modules was performed in a later step. The WGCNA protocol was directly derived from the WGCNA tutorial.

The selection of genes, based on strong connectivity, from modules selected in such a way shall hence be considered robust even if the mapping of the multi-factorial sample descriptions to a single factor, that is, the health phenotype score describing the health-effect observed in samples, may allow for plausible alternatives. This is another reason, besides the need for abstraction to compare experiments, why we consider it advantageous to compare the module’s genes against the module’s eigengene, which is derived solely by an inspection of the expression data, and not against the health phenotype score (as done in Table 3). The manual intervention to derive the health phenotype score was solely needed to filter for health(span) associated modules (Table 3).

To filter for gene interactions, we decided to filter for the strongest 5% of adjacencies from each experiment, further constrained to modules that are associated with the health(span) phenotype score; see the Methods section for details. This experiment-dependent threshold reflects that experiments differ in the number of samples and subgroups and hence in the contrasts to separate genes by their correlations.

The authors of WGCNA suggested that their software can be used to perform network meta-studies from multiple microarray experiments in a single WGCNA setup (Langfelder *et al.*, 2013). But they clearly stated that the same module needs to be robust across experiments to directly perform WGCNA on a single joint matrix based on all expression data. For the very diverse set of experiments contributing to our analysis and their polygenic phenotype this is not necessarily expected to be the case, i.e. experiments may have their true healthspan-associated module in different sections of the transcriptome. Indeed, we did not observe any interactions to have orthologs across species. The setup presented here is pragmatic and robust, i.e. individual experiments can be removed without affecting the gene interactions determined for another experiment. Of major concern for us was that hub genes are expected to show a measurable effect on health(span) only under the conditions of those ArrayExpress/GEO experiments in which they are differentially expressed. To follow this work up with wet lab confirmations, it is hence essential to provide provenance information on how the change to the hub gene’s expression was induced, i.e. a pointer to the ArrayExpress/GEO experiment. In a joint matrix across many experiments this information would be more difficult to retrieve, which suggests not to conduct the integration of experiments directly within a single WCGNA analysis.

Furthermore, for integrating interaction data from multiple experiments, the authors of WGCNA suggested to weigh the interactions from each experiment to derive a single joint adjacency matrix and they suggested to apply a threshold on that single matrix to derive a network. Because of the heterogeneity of our experiments, we cannot tell which experiment would be more informative for health(span), compared to another, and thus could not adjust weights accordingly. By treating all experiments individually, and the null hypothesis that all experiments have the same fraction of true interactions that shall be identified by the respective highest adjacency values, we could use an experiment-tailored threshold for filtering the interactions. Therefore, we used the 95^th^ percentile of correlation values in the adjacency matrix, for each experiment, to adapt the selection of the interactions to be forwarded to describe a meta-study consensus (see Figures 1 and 2). These gene interactions may be trusted and they thus could be reassembled into a larger integrated meta-study network to reflect the molecular neighborhoods of hub genes, which we presented as Figure 2 (cross-species) and Supplement Figures 2a–d (for multiple modules of the respective same species). The comparison of findings across species further strengthens the confidence in the WGCNA results. Thus, we identified conserved candidate regulators of health(span).

An important technical concern lies with the interpretation of gene expression correlation data for RNA-seq experiments, which have an intrinsic high noise-level for low-abundant genes. We have recently shown (Struckmann *et al.*, 2020) that even for array data (that are less noisy for low-abundant genes), also the low-abundant genes have a measurable effect on a ranking of genes by Pearson correlation, and this is likely also the case for module calculations as performed here. This concern has to be borne in mind in the following interpretation of the modules in terms of biological functionality.

### Cross-species hub genes and their interactions

Most of the hub genes identified by our analysis (Table 2) have been described in a health(span)-context before. The gene set enrichment analysis with g:profiler describes the molecular roles of the cross-species hub genes (Table 2) as specifically associated with a) features of the muscle and b) actin filament-based organelles and movement (Figure 3). The worm is a model species also for muscle development because of striking similarities of its muscles to mammalian muscle tissue (Christian and Benian, 2020), and movement (locomotion) is an important phenotype in all species towards operationalizing health by quantification (Fuellen *et al*. 2019). For human, rat and mouse in Table 1, there are experiments for which samples were selectively taken from muscle tissue, but not so for the worm, which is routinely sequenced as a whole. Upon closer inspection of the enrichment results of Figure 3, we found that “actin filament-based movement” refers to a wide spectrum of processes, i.e. genes that support actin polymerisation (WIPF1), the motor protein myosin (MYL1) or the transition of endosomes into exosomes for intercellular communication (SDCBP).

The number of experiments of vertebrates and invertebrates is balanced. Apart from a lack of tissue specificity, the experiments for the worm differ from rodents and humans, in that experiments for the worm may comprise samples from different larval stages. This may ease the task to find strong correlations between genes, but specificity for aging-associated processes is likely reduced.

Inspecting the distribution of hub genes by species, we found no more than five of the 12 hub genes in worm, cf. Supplement Figure 2b, and four in human, cf. Supplement Figure 2a. The only conserved direct interaction between consensus hub genes was observed between MYL1 and ACTN3 (Figure 2). However, interactions were found multiple times for experiments of the same species, namely ABRA with VRK2, AQP11 with GSTA2 and CYLD with PCNX2 for the worm. These three interactions are shown in Supplement Figure 2b and the VRK2 gene remains directly connected with the PPP1CA hub gene also after the minimum-spanning-tree-based edge removal. VRK2 is described to have downstream effects on the consensus hub gene PPP1CA (Cossa *et al.*, 2020) via GSK3beta (Lee *et al.*, 2016). Its genetic variants are associated with a series of neurological diseases and viral infection, but also with healthspan associated sleep patterns (Dashti *et al.*, 2019). The interactions conserved in multiple species are not confirmed in STRING (Szklarczyk *et al.*, 2021) for the human, but for the worm, the consensus hub gene PPP1CA (C06A1.3) links to VRK2 (tag-191).

By interpreting the enrichments in Supplement Figure 3 we can gain more insight into how the genes we discovered may be involved in health. An example is the enrichment referring to the TYROBP pathway described in wikipathways and to the GO term Leukocyte activation (Supplement Figure 3a). Genes connecting MYL1 and SKIV2L are involved in muscular structures (Supplement Figure 3b). Tubulins (e.g. TUBG1) are known to bind to PP1, of which PPP1CA is a subunit and together these proteins regulate histone acetylation (Ding *et al.*, 2008), which is reflected by the genes connecting PPP1CA and TUBG1 (Supplement Figure 3c). Further, enhanced histone acetylation is associated with extended health and lifespan in worm (Zhang *et al.*, 2009).

The highly connected genes selected in this study differ from the list we recently published (Möller *et al.*, 2020). This WGCNA-based study does not refer to prior knowledge about genetic contributions and does not perform a factor analysis to directly associate genes with a health(span) phenotype. Instead, our focus here is the network-centric interpretation of correlations within gene co-expression clusters, i.e. WCGNA modules. It is the module as a whole that correlates in its expression with health, not necessarily the individual genes. Table 3 lists genes within the clusters that are most representative for the features/characteristics of the cluster in question, i.e. that have the highest degree of *module membership* by WGCNA definition, and in the table, there are marks (by boldface) for the subset of genes that are also hub genes. Of the cross-species hub genes in Table 2, six are also listed in Table 3. Others are “near misses”, e.g. Table 3 does not list the consensus hub gene MRPL18 but MRPL19. And besides the consensus hub gene PPP1CA, other PP1 subunits like PPP1CB and PPP2R3C are found in two species (Table 3). PPP1CB was also found as a hub gene, but only for the worm.

In Table 3, we report SQSTM1 as the only gene that is associated with health in three species. That gene was long suggested to be aging- and health-related (Bitto *et al.*, 2014; Sánchez-Martín and Komatsu, 2018), also for human, even though it was only found associated in the analyses of the animal experiments in this study. Its transcript is negatively correlated with health, but SQSTM1 overexpression is known to extend healthspan in worm (Kumsta *et al.*, 2019), which may be suggestive for a protective upregulation effect.

Overall, our meta-analysis of a very diverse set of transcriptomics experiments successfully identified genes which, for the most part, were already established to be closely associated with health(span), and together they have a strong and meaningful GO term enrichment. The enrichment of muscle-related genes can be credited to our focus on health(span) experiments, and our study found many “actin filament-based movement” genes (Figure 3) that provide the cellular infrastructure not just for movement, but also for signalling and cell division, which may be triggered/blocked whenever cells start to feel unwell. If so, then it may be possible to detect many healthspan genes solely by inspecting cellular data. This hypothesis may be confirmed by an extension of our setup to a larger set of cellular transcriptomics data sets for which samples vary in their genetic or environmental exposure to stress factors.

This study provided a cross-species meta-study of gene interactions for health(span)-related datasets in ArrayExpress/GEO. It focused on a co-expression network analysis and subsequently on derived hub genes, instead of a focus on those genes that correlate the most with the healthspan phenotype score. This approach shall allow for an abstraction from the experiment at hand and permit a search for common mediators of an effect. The proposed consensus hub genes were plausible in their implication into health(span). Their interactions could be confirmed in STRING, or were found consistent with gene set enrichment analyses and they may support the interpretation of joint or epistatic effects between pairs of haplotypes in healthspan GWAS or linkage analyses. The protocol as provided with WGCNA is very transparent so that findings can be traced back to the experiments that are backing them, to serve as a template for further investigations in the wet lab.

## Acknowledgements

This work was supported by a project from the European Union’s Horizon 2020 research and innovation programme under Grant agreement No 633589 (Aging with Elegans), and by the BMBF (Validierung des technologischen und gesellschaftlichen Innovationspotenzials wissenschaftlicher Forschung – VIP+, FKZ: 03VP06230).

## Availability

Our implementation is available online at https://bitbucket.org/ibima/healthspantranscriptomicsnetworks/.

---

**Supplement Figure 1a:**
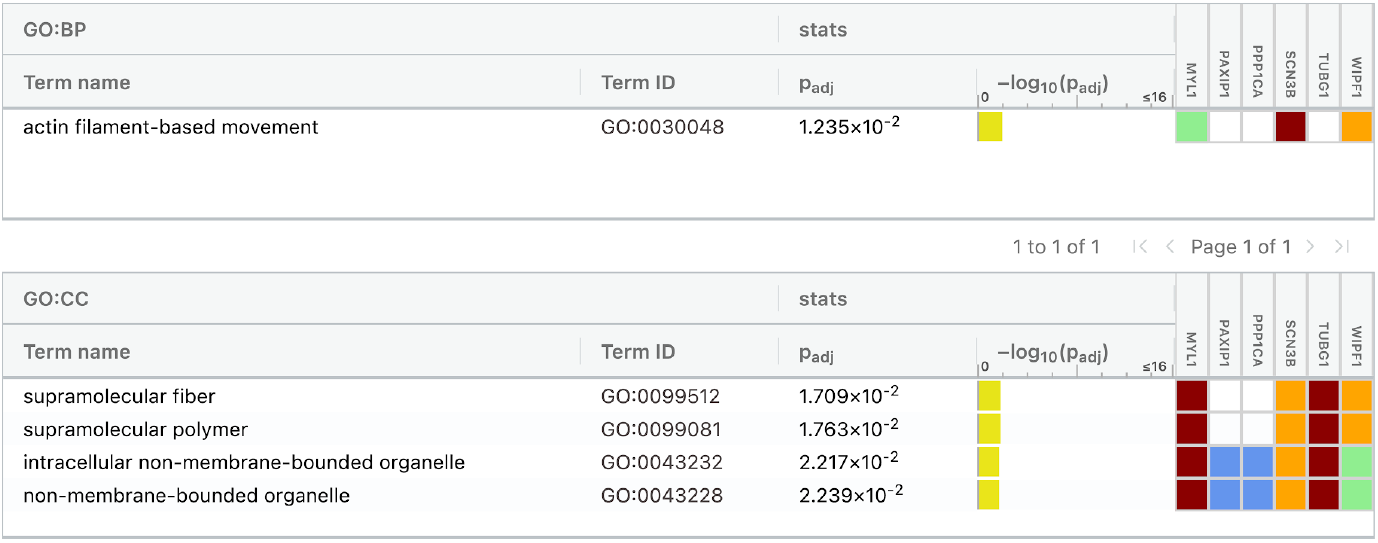
g:profiler gene set enrichment analysis of genes listed jointly in Tables 2 and 3.

**Supplement Figure 1b:**
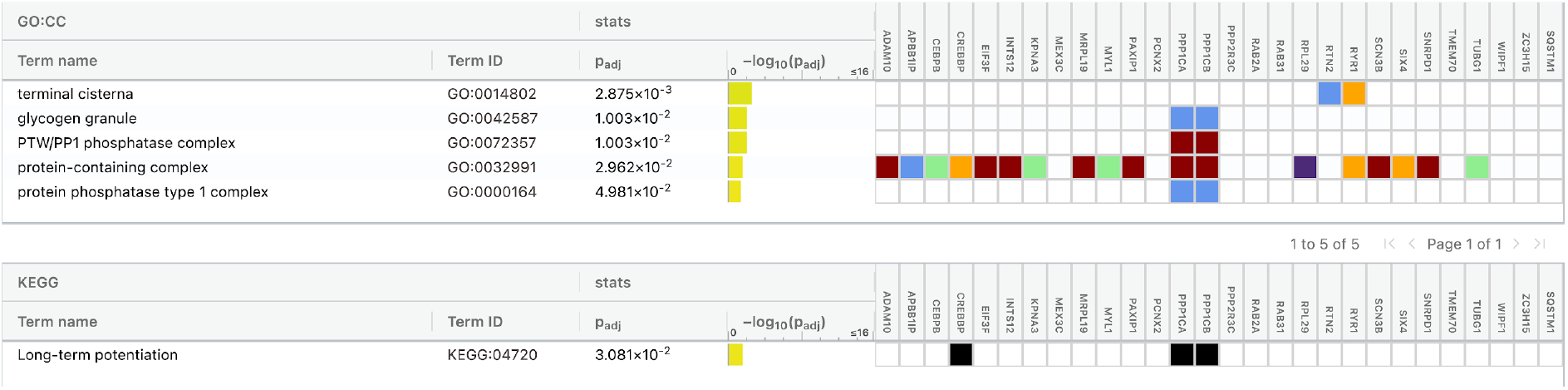
g:profiler gene set enrichment analysis of genes listed in Table 3.

**Supplement Figure 2a:**
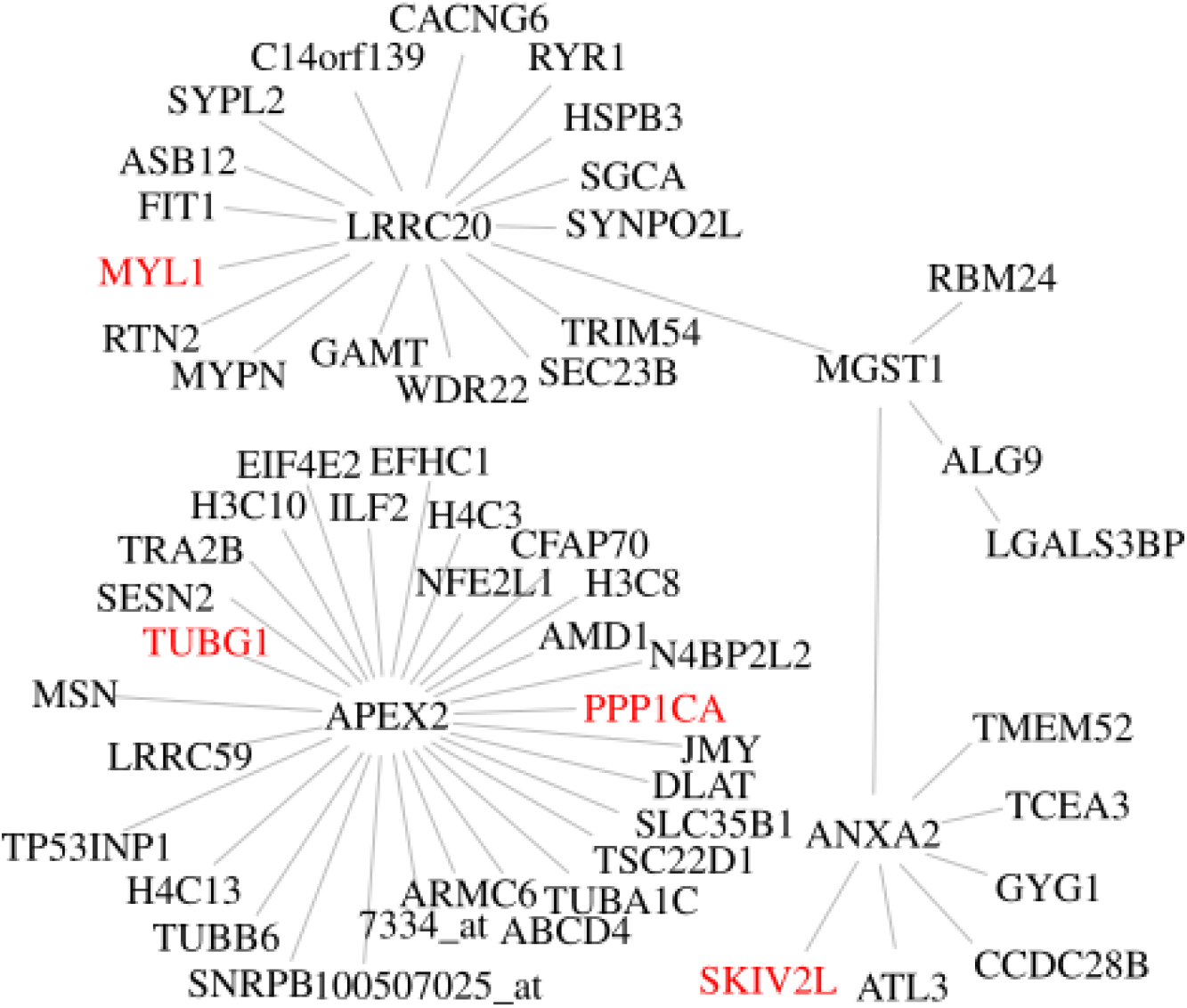
Gene interactions observed in human. The genes MYL1, PPP1CA, SKIV2L and TUBG1 are hub genes in WGCNA-defined modules from multiple species (cf. Figure 1, here shown in red). This graph was created iteratively with the red genes as a seed, then adding all the gene-gene interactions from WGCNA in human experiments originating from these red genes, and then transitively adding all the interacting genes of those. The resulting graph is highly interconnected before applying the minimum spanning tree algorithm. The genes interacting with hub genes across species (in red) then appear marginalized by the three human-only hub genes ANXA2, APEX2, LRRC20 and MGST1, given that the minimum spanning tree shows only the strongest correlations. ANXA2 is well described for a wide array of disease, i.e. cancer but also pulmonary fibrosis, and on a molecular level chimes in with vesicle fusion. APEX2 is a nuclease required for lymphocyte proliferation. LRRC20 is not yet described but known to interact with the also mostly undescribed TOM1 that once more is thought to be involved in intracellular trafficking and the E3 SUMO-protein ligase ZBED1. MGST1 is an enzyme located at the ER and mitochondria, a transferase of glutathione, an antioxidant.

**Supplement Figure 2b:**
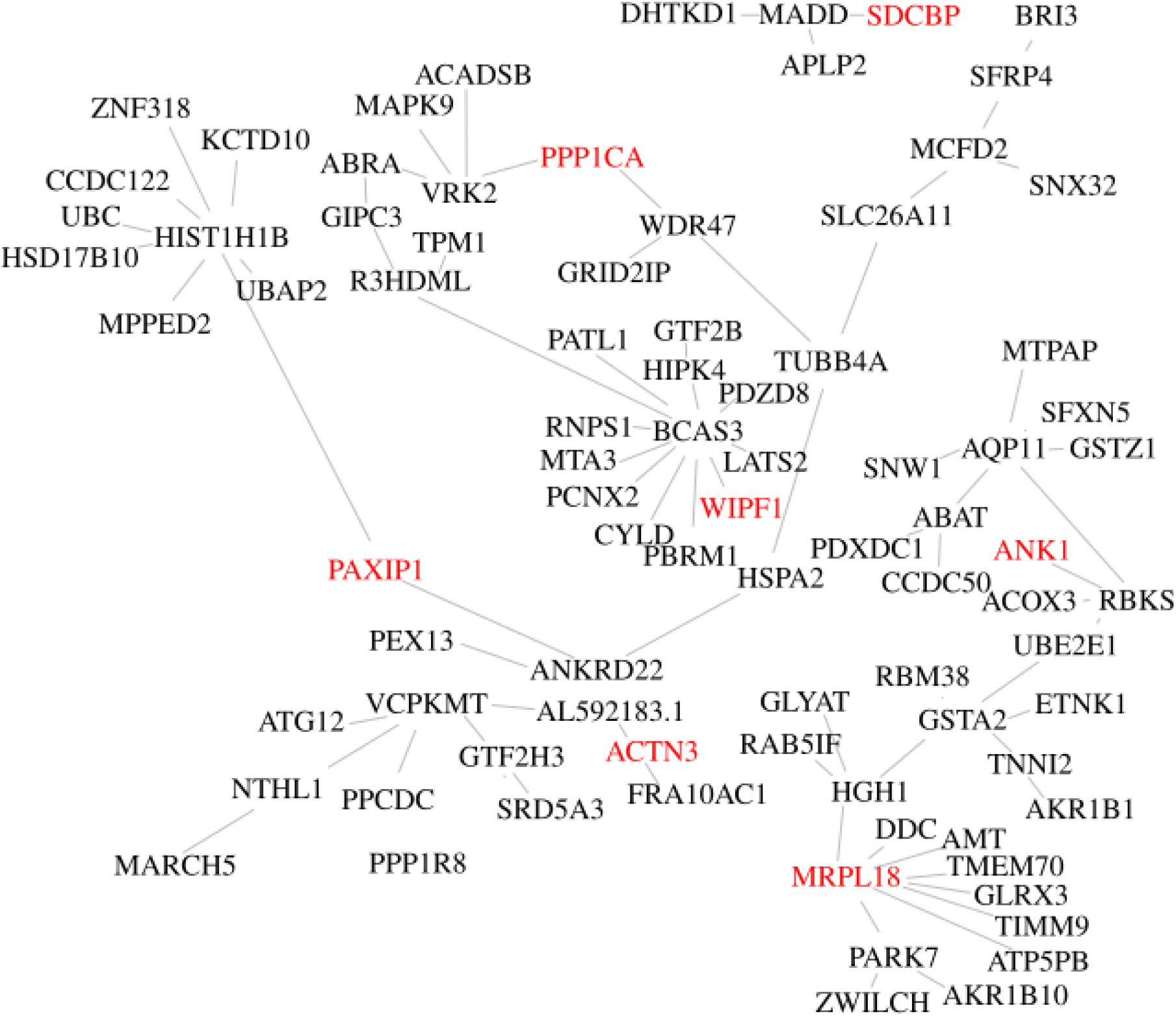
Gene interactions observed in worm. The figure was prepared analogously to Supplement Figure 2a from gene interactions observed in the worm. Gene names were mapped to human orthologs for an easier comparison between species.

**Supplement Figure 2c:**
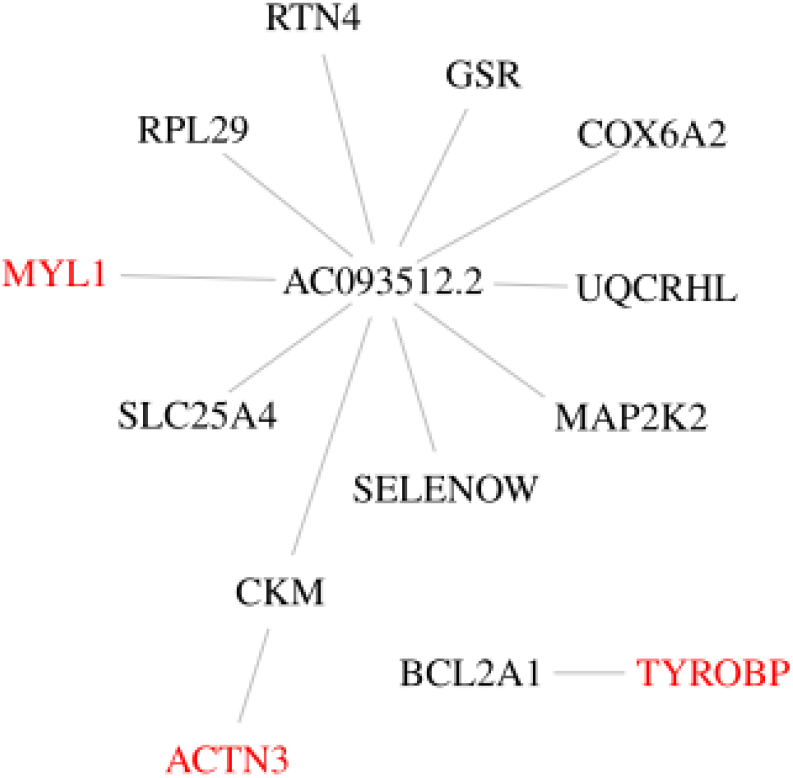
Gene interactions observed in rat.

**Supplement Figure 2d:**
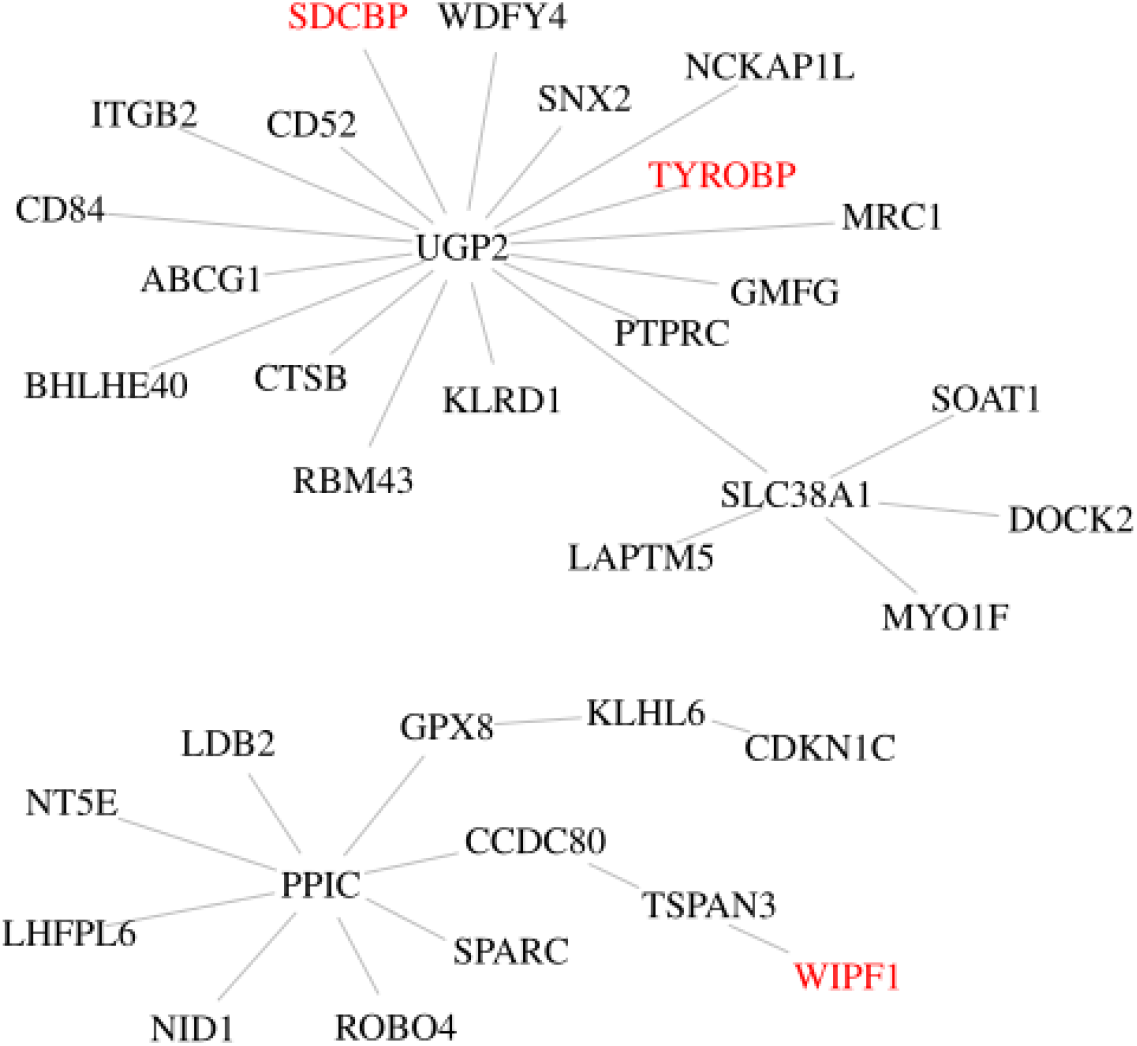
Gene interactions observed in mouse.

**Supplement Figure 3a:**
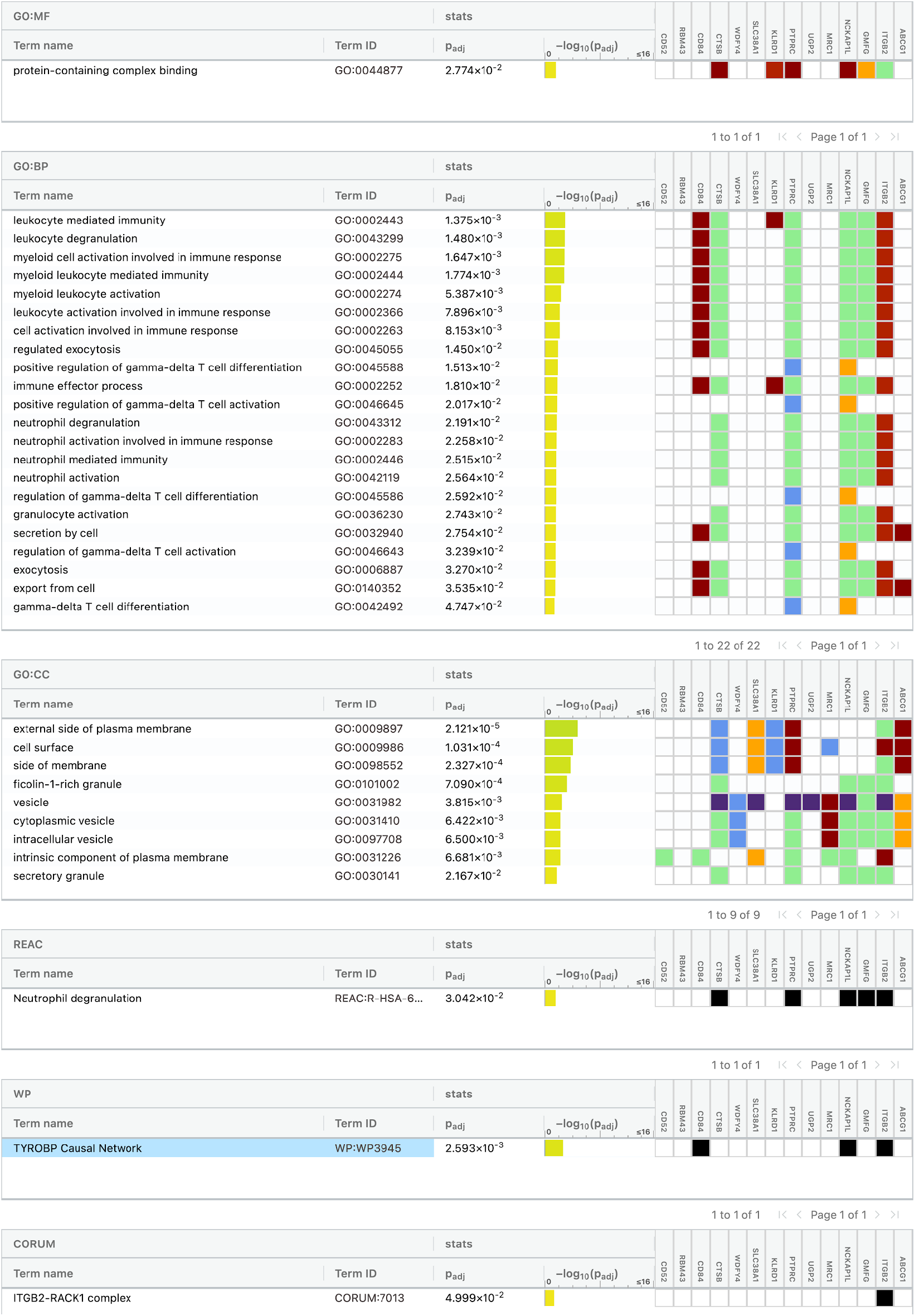
g:Profiler GSEA on genes connecting hub genes TYROBP and SDCBP.

**Supplement Figure 3b:**
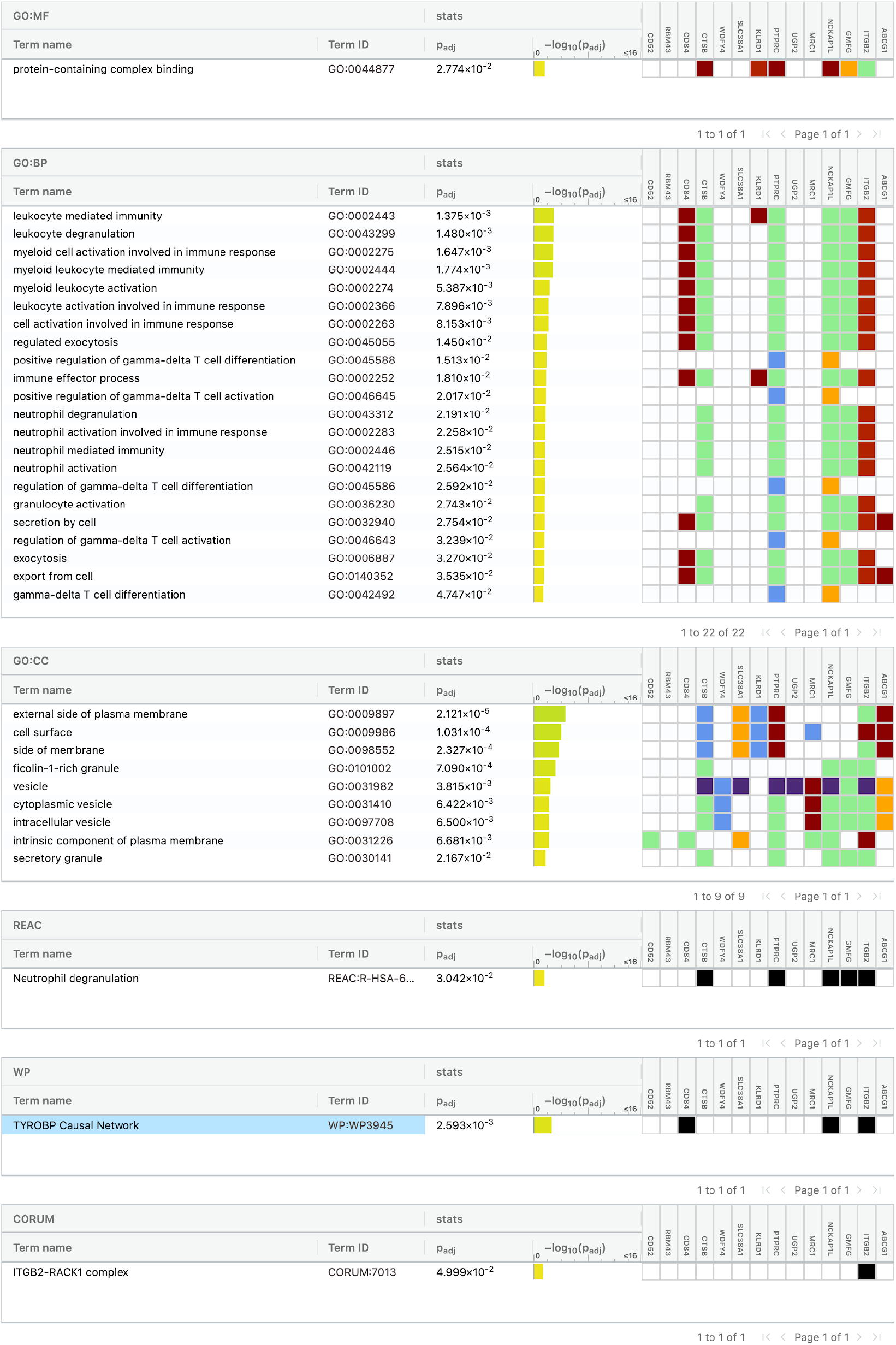
g:Profiler GSEA on genes connecting hub genes MYL1 and SKIV2L.

**Supplement Figure 3c:**
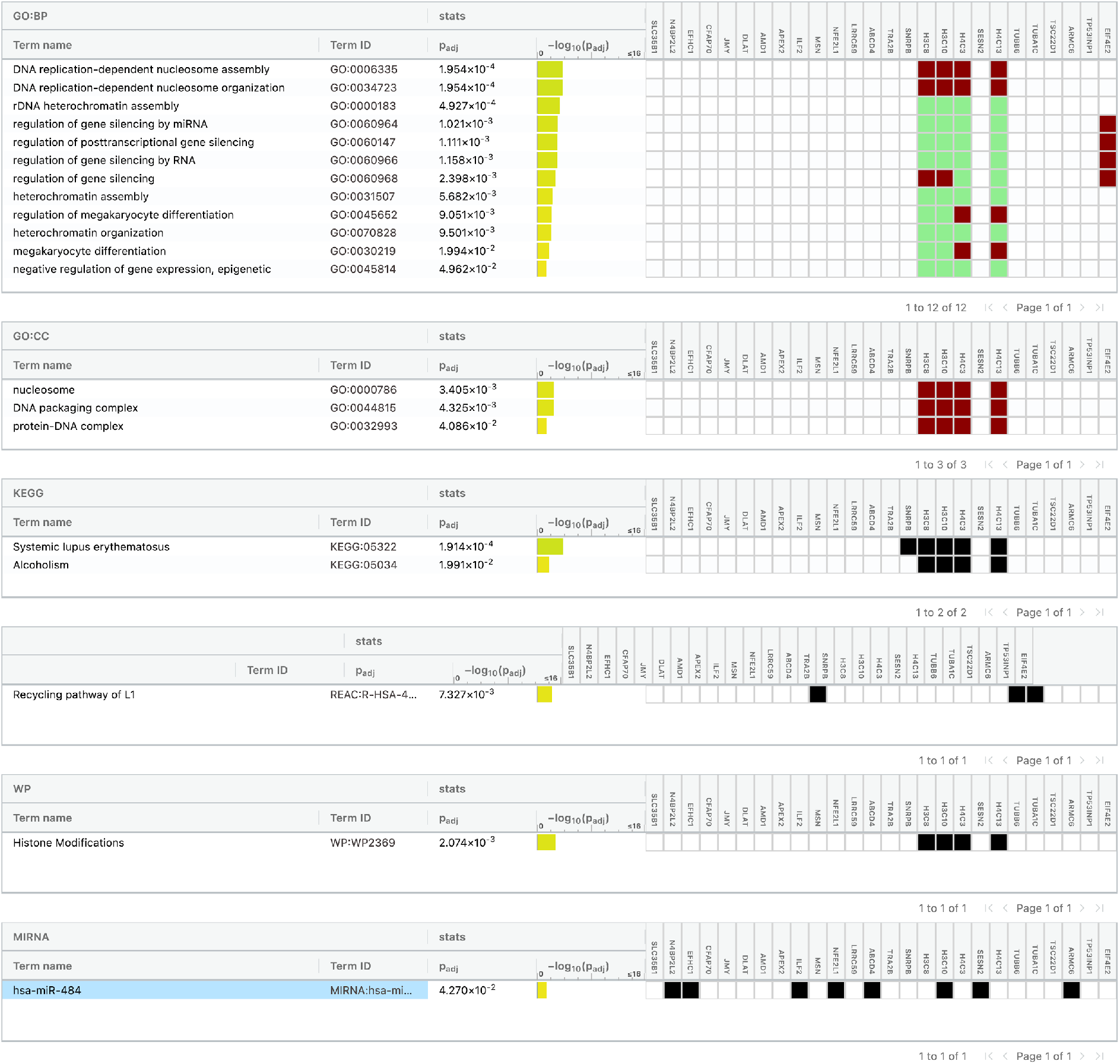
g:Profiler GSEA on genes connecting hub genes TUBG1 and PPP1CA.

**Supplement Table 1:**
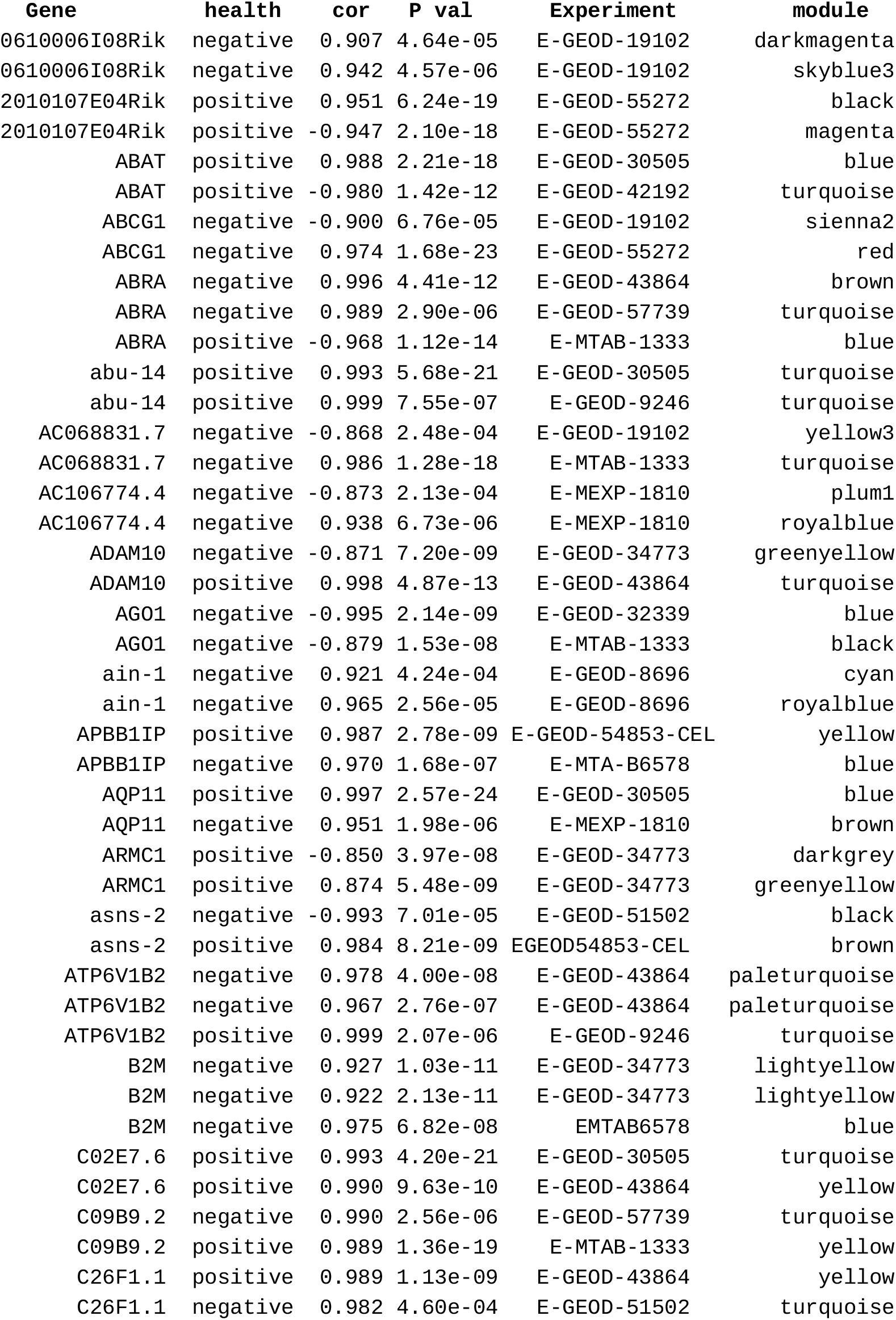
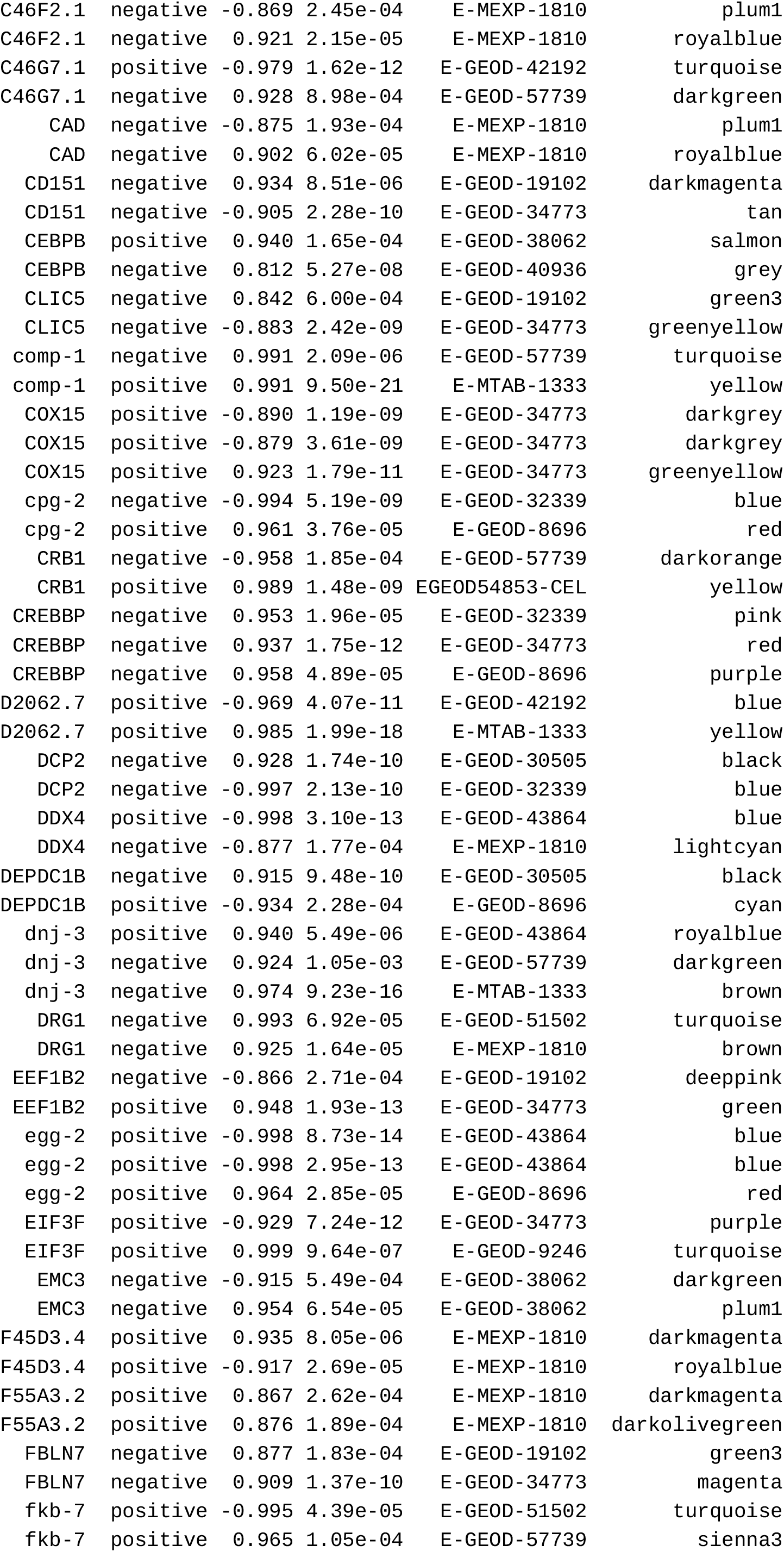
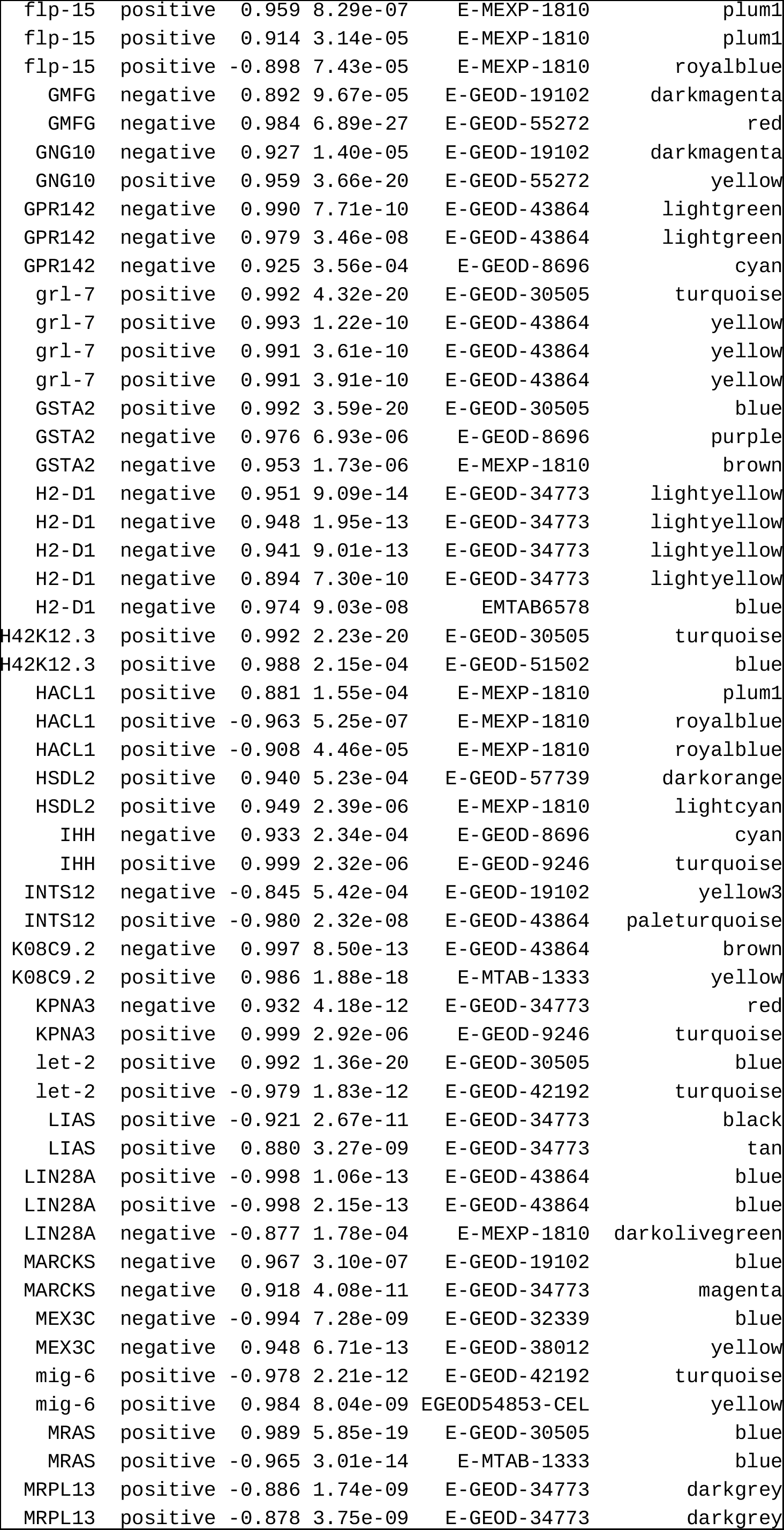
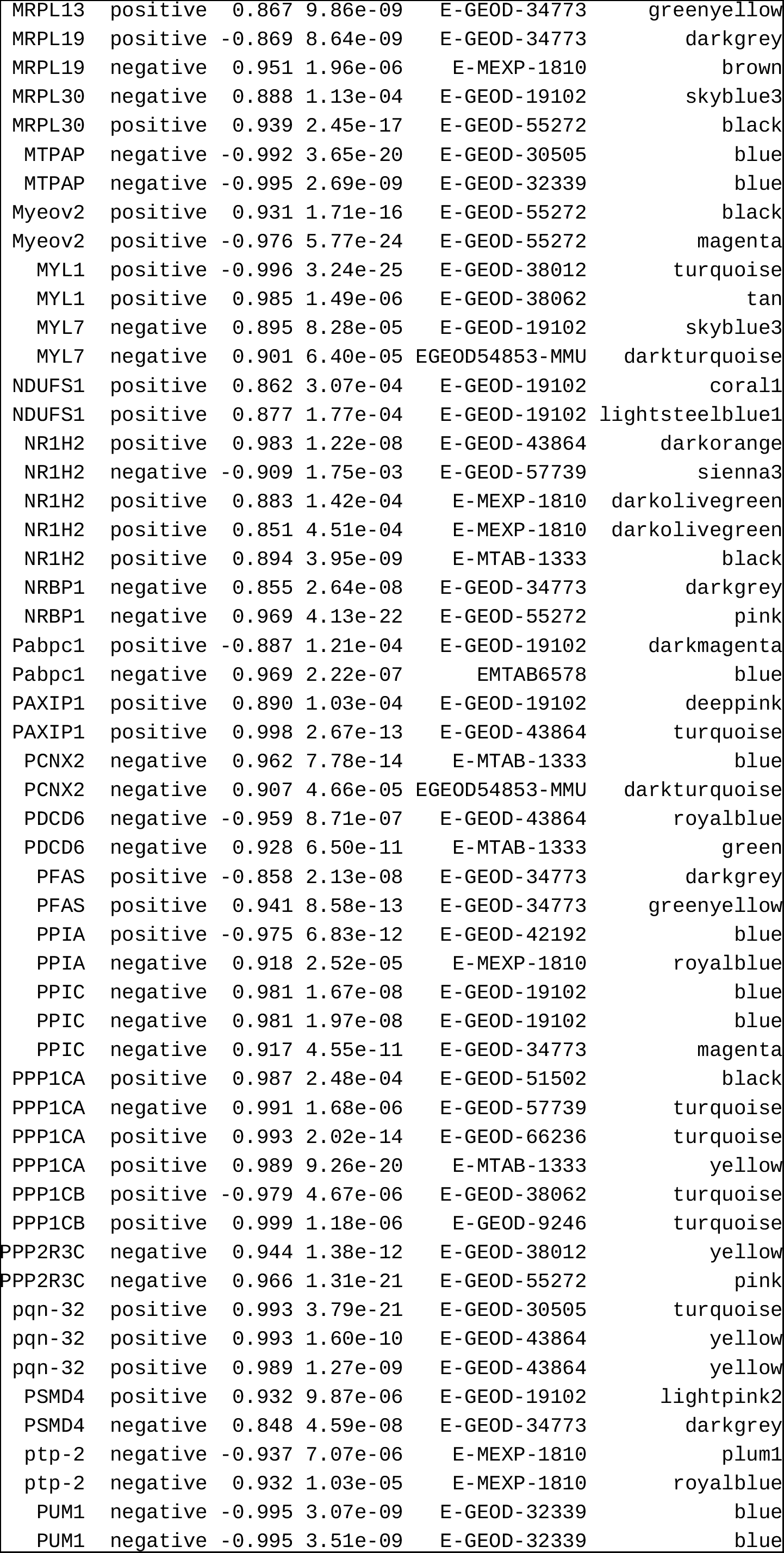
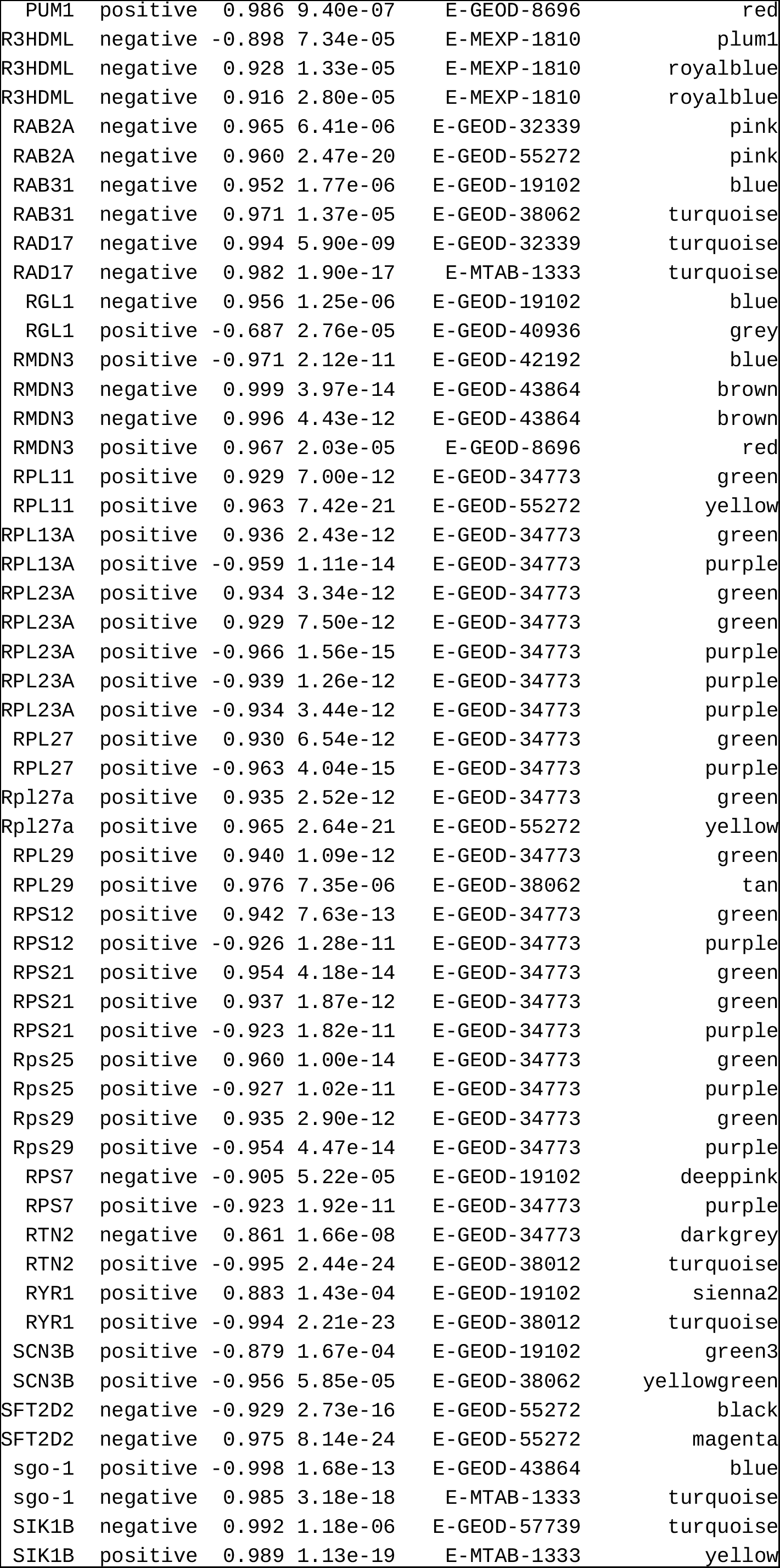
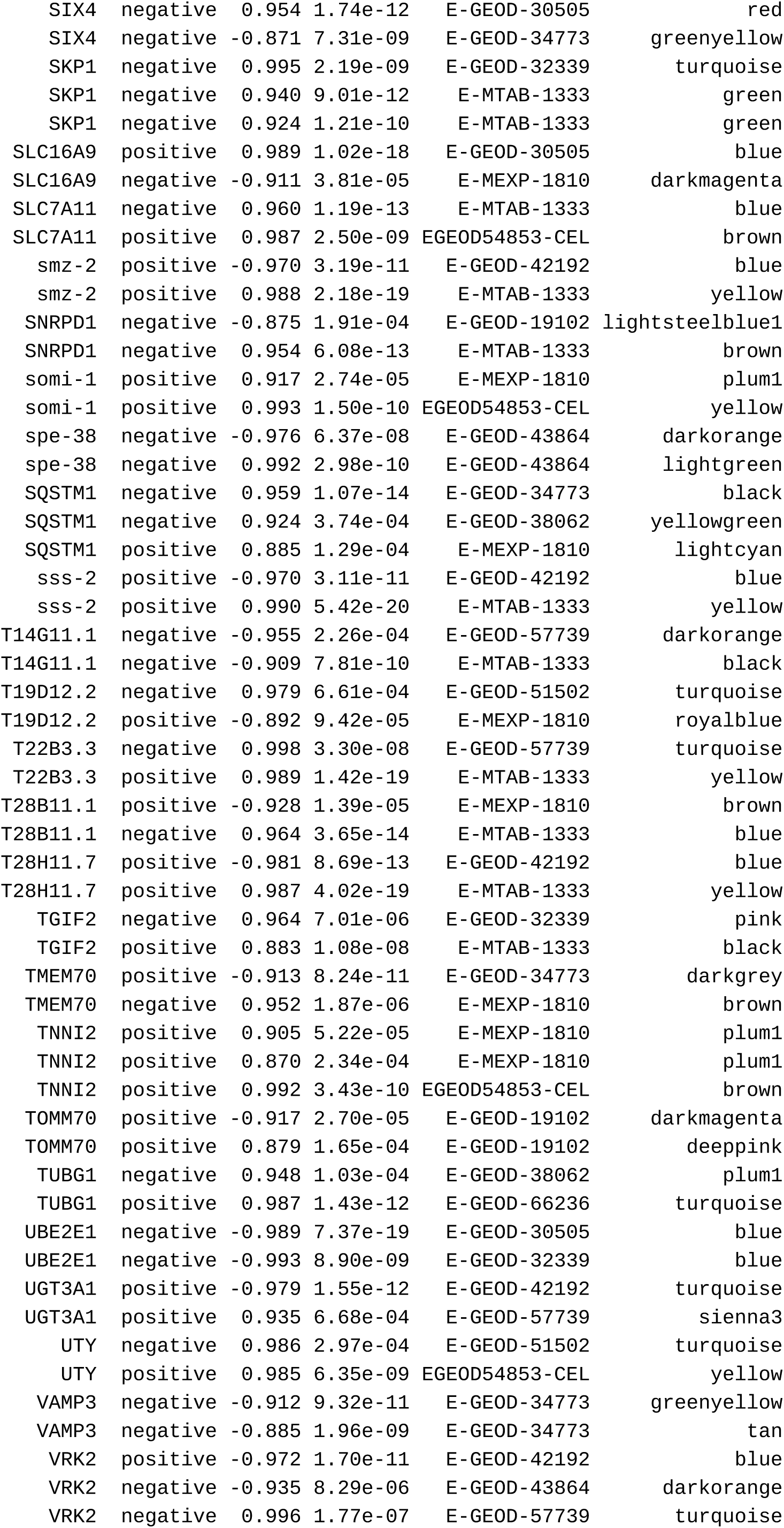
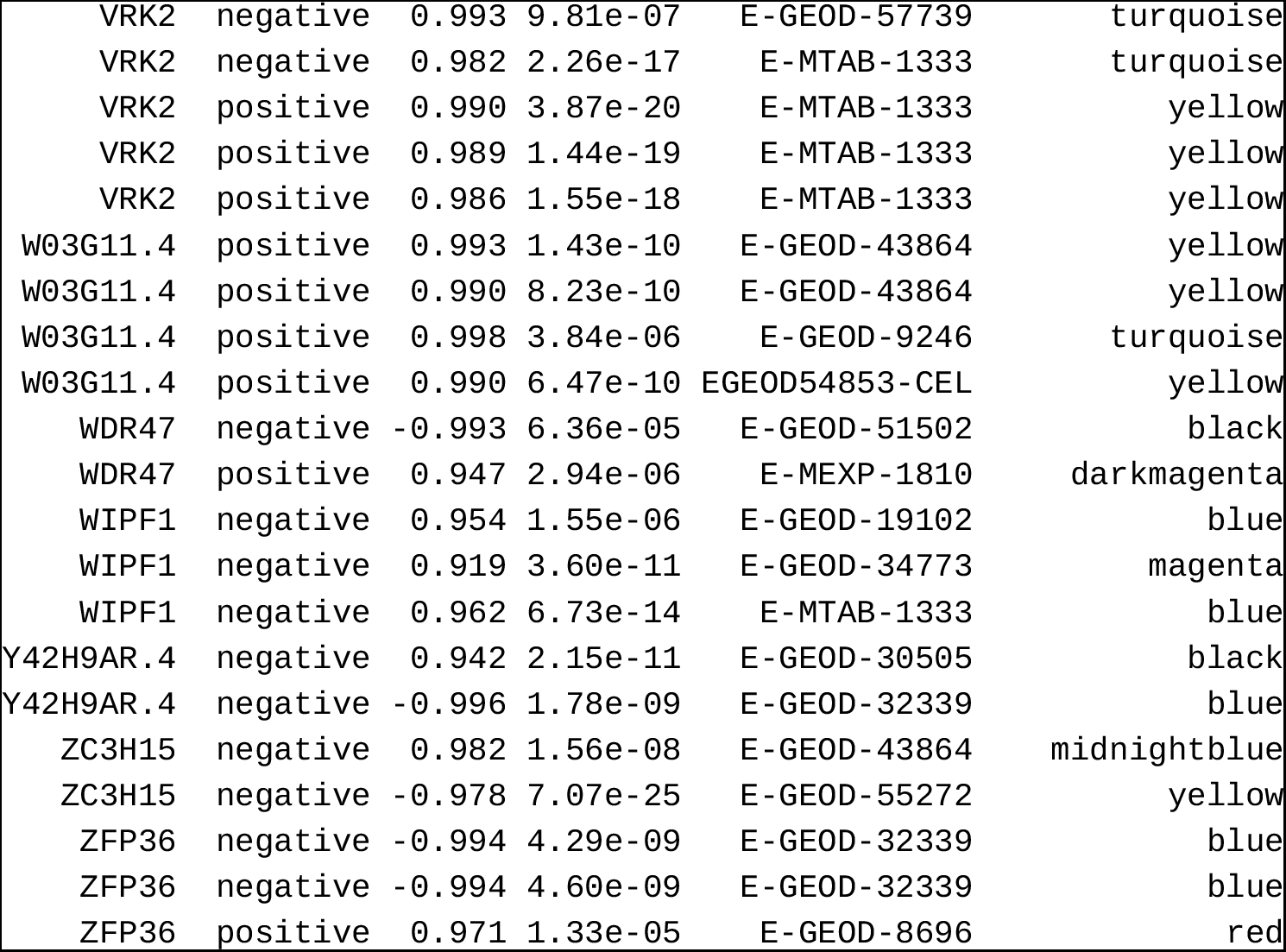
Genes correlating with eigengene representation (module membership) of those modules that are correlating with the health score. The table lists all genes that appear in at least two experiments among the top 30 and a P value below 0.05. If the expression of the gene correlates positively with the health score then the gene is tagged as positive. The columns “cor” and “P val” list the values as determined by WGCNA. A negative correlation with a positive tag indicates that eigengene of the module is negatively correlated with the health score.

